# Regulation of Polyhomeotic condensates by intrinsically disordered sequences that affect chromatin binding

**DOI:** 10.1101/2021.10.04.463094

**Authors:** Ibani Kapur, Elodie L. Boulier, Nicole J. Francis

## Abstract

The Polycomb group (PcG) complex PRC1 localizes in the nucleus in the form of condensed structures called Polycomb bodies. The PRC1 subunit Polyhomeotic (Ph) contains an oligomerizing sterile alpha motif (SAM) that is implicated in both PcG body formation and chromatin organization in *Drosophila* and mammalian cells. A truncated version of Ph containing the SAM (mini-Ph), forms phase separated condensates with DNA or chromatin *in vitro*, suggesting PcG bodies may form by SAM-driven phase separation. In cells, Ph forms multiple small condensates, while mini-Ph typically forms a single large nuclear condensate. We therefore hypothesize that sequences outside of mini-Ph, which are predicted to be intrinsically disordered, are required for proper condensate formation. We identified three distinct low complexity regions in Ph based on sequence composition. We systematically tested the role of each of these sequences in Ph condensates using live imaging of transfected *Drosophila* S2 cells. Each sequence uniquely affects Ph SAM-dependent condensate size, number, and morphology, but the most dramatic effects occur when the central, glutamine rich IDR is removed, which results in large Ph condensates. Like mini-Ph condensates, these condensates exclude chromatin. Chromatin fractionation experiments indicate that removal of the glutamine rich IDR reduces chromatin binding, while removal of either of the other IDRs increases chromatin binding. Our data suggest all three IDRs, and functional interactions among them, regulate Ph condensate size and number. Our results can be explained by a model in which tight chromatin binding by Ph IDRs antagonizes Ph SAM driven phase separation and highlight the complexity of regulation of biological condensates housed in single proteins.

## INTRODUCTION

The role of biomolecular condensates in regulation of cellular processes, including gene expression, is increasingly appreciated. Phase separation, the demixing of molecules in solution to dense and dilute phases, may underlie many condensates that form in cells, providing a means to concentrate and segregate biomolecules [1–3]. However, while the core physical principle of phase separation may explain condensates, phase separation must be tightly regulated in time and space to generate condensates with biological functions. The material properties of condensates, which can include more or less viscous liquids and gel-like or even solid states, are also essential for their function [4, 5]. Finally, biological condensates are not simple polymer in solvent systems, but highly complex structures existing in an equally complex solvent (i.e. the cytoplasm or nucleus) [6].

To understand how complex condensates form in cells, identifying key proteins that drive their formation, and determining how the sequences of these proteins contribute to condensates is essential. Many examples, as well as theory, have demonstrated how both structured domains and intrinsically disordered sequences can undergo phase separation under biologically relevant conditions [7]. Because weak, multivalent interactions are key to creating liquid condensates, Intrinsically Disordered Regions (IDRs) are especially well suited to undergo phase separation [1, 7, 8]. However, IDRs may lack the specificity needed to form relevant condensates in the complex cellular environment [9]. There are now several examples of proteins that drive condensate formation through a combination of structured domains and IDRs [10, 11].

The Polycomb Group (PcG) proteins are epigenetic regulators of gene expression that function by modifying chromatin structure [12–14]. Two main PcG multiprotein complexes, PRC1 and PRC2, are conserved across evolution [15]. Both complexes modify histones (H2A119Ub and H3K27me3, respectively). PcG proteins also organize chromatin at a large scale, forming large, compacted chromatin domains [13, 16]. This activity is distinct from histone modifications. PcG proteins themselves form visible condensates, and these condensates co-localize with PcG-regulated genes [13, 16]. This suggests that the formation of PcG condensates and of compacted chromatin domains are linked processes. PRC1 components, particularly Polyhomeotic (Ph) and Polycomb (Pc) (PHC and CBX in mammals) are the most implicated in condensate formation and chromatin organization [17, 18].

Ph has a C-terminal Sterile Alpha Motif (SAM) that is essential for its function and implicated in both condensate formation and chromatin organization. The Ph SAM can form head to tail polymers [19], and this polymerization activity is important for Ph function [20]. Dominant negative Ph SAM mutations that disrupt polymerization of wild-type Ph SAM also disrupt PcG clusters, lead to loss of chromatin compaction, and disrupt gene regulation in both *Drosophila* and mammalian cells, and in mice [21, 22]. Loss of Ph leads to decompaction of chromatin in *Drosophila* cells, *Drosophila* embryos, and mouse embryonic stem cells [17, 23, 24]. Although SAM polymerization is its hallmark activity, genetic experiments indicate that Ph SAM has critical polymerization-independent functions because a *ph* transgene with a polymerization interface mutated can rescue some *ph* functions in *Drosophila* embryos, while *phΔSAM* cannot [25].

We recently showed that Ph SAM, in the context of a truncated version of Ph containing its three conserved domains, called mini-Ph, undergoes phase separation with DNA or chromatin *in vitro*. Phase separation requires the SAM, but does not strictly require polymerization [26]. Human PHC1 was also shown to undergo SAM-dependent phase separation in cells when induced to cluster through optogenetic manipulation [27]. Thus, phase separation could be the critical second Ph SAM activity. However, in cells, Ph forms many condensates, while mini-Ph forms a single condensate; both types of condensates require the SAM [26]. Thus, we hypothesized that sequences N-terminal to mini-Ph regulate condensate formation through the SAM. Indeed, *Drosophila* Ph has ∼1200 additional amino acids N-terminal to mini-Ph that have no recognizable domains and are not characterized. Most mammalian PHC isoforms (all of which have the mini-Ph region) also contain uncharacterized sequence N-terminal to the SAM.

Here, we analyze the N-terminal sequence of Ph, which is predicted to be intrinsically disordered. We divided the N-terminal sequence into three distinct IDRs based on sequence composition and complexity, and tested their effects on condensate formation in cells. We find that all three IDRs influence condensate formation. The most prominent effects are mediated by a central, glutamine-rich IDR; deletion of this region results in large, round condensates that exclude chromatin, similar to those formed by mini-Ph. Deletion of this IDR also strongly reduces chromatin association, suggesting the effects of this sequence on condensate properties are determined by chromatin binding. The other IDRs also affect condensate properties and chromatin binding. This includes an IDR previously shown to regulate Ph SAM aggregation through its O-linked glycosylation, which we find also negatively regulates the activity of the other two IDRs. Our results point to chromatin binding as an important constraint on phase separation in cells, and suggest balance between the two has evolved to tune biological condensates.

## RESULTS

### The N-terminal region of Drosophila Ph is predicted to be disordered and comprises three distinct regions

Drosophila melanogaster contains two tandem ph genes (ph-p and ph-d), which are highly similar and largely functionally redundant[28, 29]. Our analysis is of Ph-p, referred to hereafter as Ph. Ph is a large protein (1589 amino acids), with three small conserved domains in its C-terminal region namely the HD1, FCS and SAM (**Figure 1A**). Previous work noted biased sequence composition in the region N-terminal to the conserved domains[30], and characterized an unstructured S-T rich region that is both phosphorylated and heavily glycosylated by the O-GlcNac transferase (OGT), encoded by the PcG gene super sex combs (sxc) [25, 31]. We used MetapredictV2 [32, 33], which integrates disorder prediction and AlphaFold2 structure prediction, to analyze the sequence of Ph. This algorithm classifies the entire N-terminal region of Ph upstream of the SAM as disordered (**Figure 1A**). This includes the FCS and HD1 domains, both of which have solved structures from human Ph homologues (PHC2 and PHC1, respectively) [34, 35]. Although classified as disordered, these regions show a dip in disorder and corresponding increase in structure. The structure of the HD1 was obtained in the presence of a (fused) binding partner; it therefore seems possible that this is an induced fold, and thus that the sequence is disordered in an isolated polypeptide. We used MetapredictV2 to analyze all three PHC sequences; in all cases, the FCS, but not the HD1, is predicted to be ordered **(Figure S1)**, indicating that differences in the FCS sequence explain the difference in predicted structure. A small possibly structured region is predicted around aa400 of Ph, and a larger region that corresponds to a Q-rich stretch (grey bars, **Figure 1A**). This region is predicted to form a long helix by AlphaFold2, and also to form coiled-coils (not shown). As discussed recently, predictions of coiled coil and helical structures from poly-Q regions are not well supported by structural data [36], although the importance of helix forming ability of a PolyQ sequence in regulating phase separation has also been demonstrated [37]. Like any prediction, the disordered nature of the N-terminal sequences in Ph will need to be confirmed experimentally; we refer to the entire region as disordered, with the above caveats noted (**Figure 1A**).

**Figure 1.**
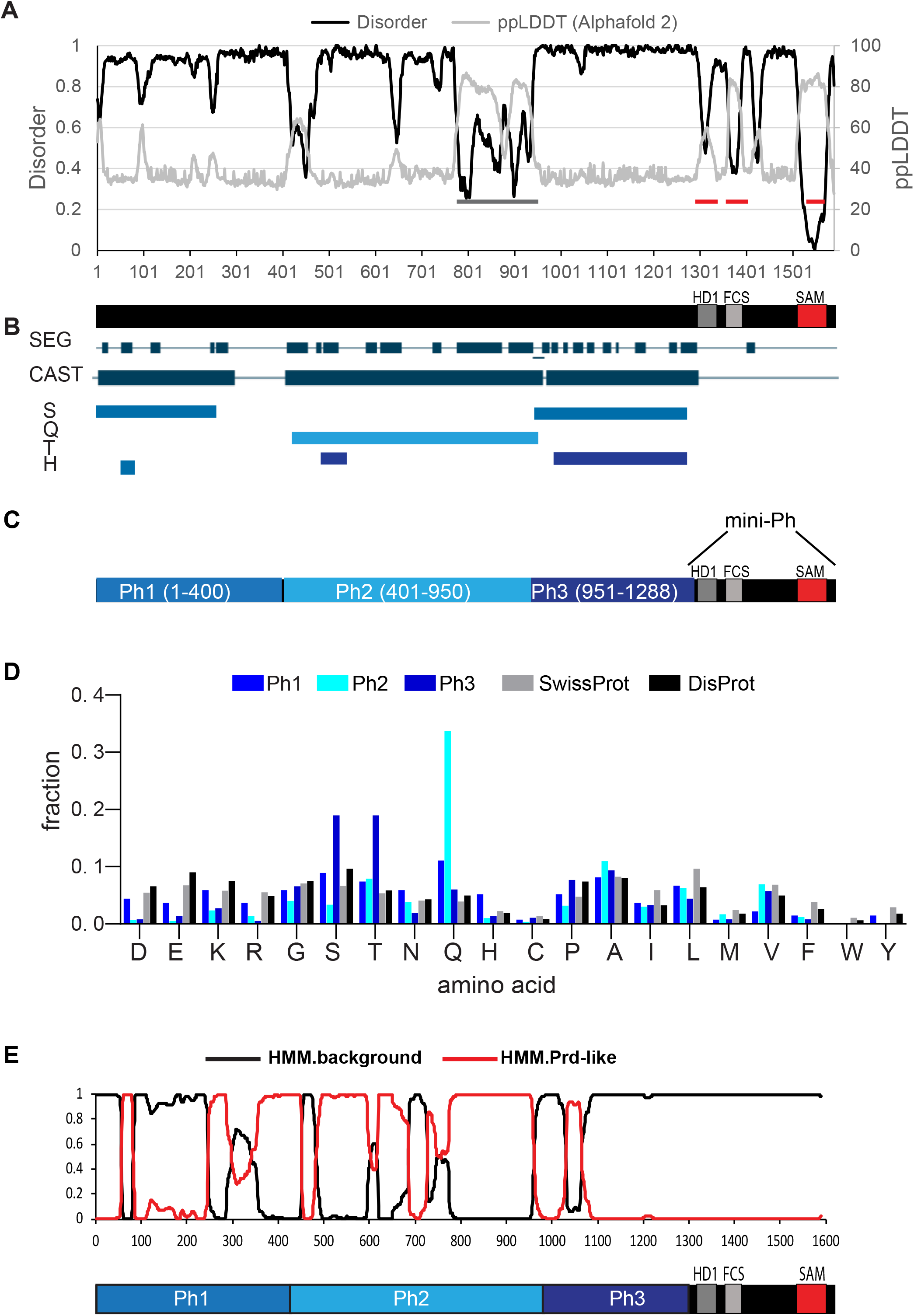
Identification of three IDRs in Ph. (**A**) Prediction of intrinsically disordered sequence using MetapredictV2, which integrates AlphaFold 2 predictions[32], [33]. X-axis is amino acid position. Most of Ph is predicted to be disordered (>0.5 on left Y-axis), with the exception of the Ph SAM (indicated with a red line). The two other red lines correspond to the HD1 and FCS motifs, which are scored as disordered, although the AlphaFold2 signal predicts some order. The grey bar indicates a region predicted to have some order; this is a densely Q-rich region that is predicted to form helices/coiled-coils. (**B**) SEG/CAST analysis of Ph using PlaToLoCo [39] suggests the disordered region can be parsed into subregions based on which amino acids are masked (repetitive). CAST track indicates all masked regions, and the masked amino acids are indicated below. (**C**) Schematic of different regions in the Ph disordered region defined as Ph1(IDR1), Ph2(IDR2) and Ph3(IDR3). (**D**) Frequency of each amino acid in the three IDRs, relative to the SwissProt and DisProt data bases. (**E**) Prion like domain prediction using PLAAC[40]. IDR 2 is dominated by high scoring prion-like regions, but all three IDRs have predicted PrDs. See **sFig. 2** for full sequence of each IDR.

Intrinsically disordered protein sequences often also contain low complexity and compositionally biased sequence. To identify such regions in Ph, and potential subregions in the large disordered region, we used SEG and CAST [38] algorithm using the PlaToLoCo (Platform of Tools for Low Complexity) interface [39]. Although low complexity sequence is present throughout the N-terminal region, the masked (i.e. repetitive) amino acids are different in different regions of the sequence. This allowed us to demarcate Ph1 (S-masked); Ph2 (Q-masked); and Ph3 (S+T masked) (**Figure 1B**). Ph1 does not have a clear enrichment for a single amino acid, but Ph2 is enriched in Glutamine (36%) and Ph3 in Threonine/Serine (20% each) (**Figure 1C, D**). PLAAC was used to scan for prion like amino acid composition in the protein sequence [40]. Consistent with its high glutamine content, Ph2 has high scoring predicted prion-like regions; both Ph1 and Ph3 have smaller a predicted prion-like regions (**Figure 1E**). In summary, we refer to the three Ph regions as Ph(IDR)1-3, and have used these boundaries to design constructs to test their functions. The full sequence of each IDR is provided in **Figure S2**. Although the sequence composition rationalizes the boundaries chosen, they must be regarded as somewhat arbitrary. The boundaries split the disordered region into three roughly equal sized pieces making them a good starting point for structure function analysis.

### Expression of Ph proteins lacking each IDR, or combinations of them, in cells

To understand the contribution of each IDR to Ph condensate formation, we designed a series of constructs consisting of N-terminal fusions of the Venus GFP variant to truncated versions of Ph. Constructs were designed to test: 1) the effect of removing each IDR from Ph; 2) the activity of each IDR when fused to mini-Ph (i.e. deletion of 2 IDRs); 3) the role of the SAM in IDR effects; 4) the condensate forming activity of each IDR alone or in combination. We used the heat shock promoter for inducible expression. Constructs were transiently transfected into Drosophila S2 cells, along with a plasmid encoding H2Av-RFP as a nuclear marker. Cells were subjected to mild heat shock (8 min. at 37°C) to induce protein expression, and allowed to recover overnight before live imaging.

We first used Western blots to confirm that full-length protein is expressed for each construct, and that the expression levels are similar. There is ∼3x variability in expression levels for different proteins, although transfection conditions were identical (**Figure S3A-D**). We also compared expression with endogenous Ph for proteins that maintain the antigen recognized by our antisera (**Figure S3E-F**). Average total Ph levels did not exceed 2-fold of untransfected cells, due to downregulation of endogenous ph. The Ph locus is known to contain Polycomb Response Elements [41], which may mediate auto-regulation [42, 43].

### Analysis of the function of Ph IDRs using live imaging

As shown previously [26], transfected wild type (WT) Ph forms several round, bright condensates in cells, while mini-Ph forms no condensates or a single condensate (**Figure 2A-E**). We carried out two analyses on the imaging data (see Methods for details). To analyze large numbers of cells in an unbiased manner, we developed Cell Profiler [44, 45] analysis pipelines to identify condensates, count them, and measure their size. We also carried out a smaller manual analysis using ImageJ, to count condensates and measure total nuclear intensity of Venus. Imaging conditions were held constant; however, because there are a range of expression levels in transfected cells, and because we aimed to detect both small and large condensates, images of some cells were clearly saturated. This will result in underestimation of intensities, but should only affect the highest expressing cells.

**Figure 2.**
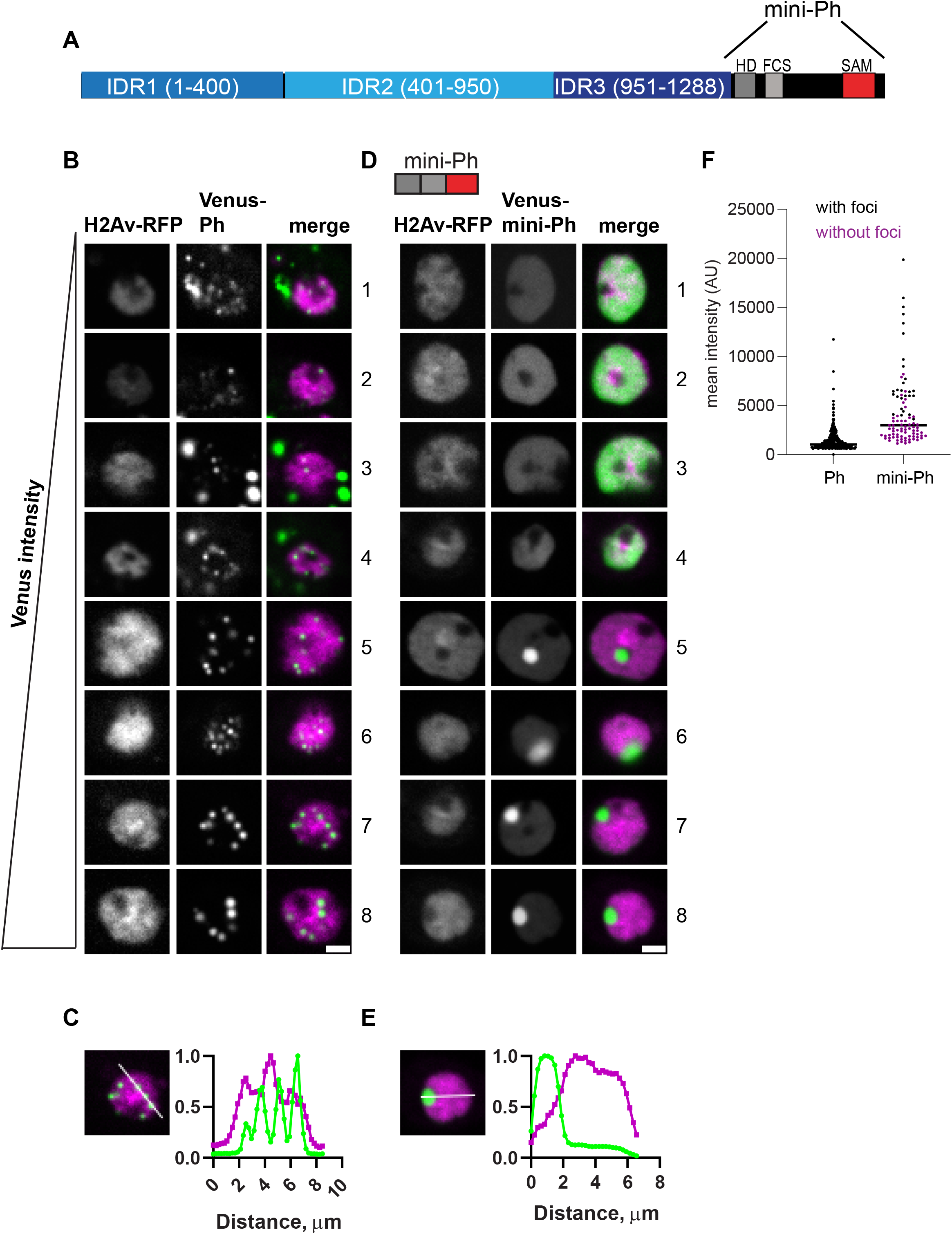
Characterization of Ph and mini-Ph condensates and their relationship to protein expression level. **(A)** Schematic of Polyhomeotic. **(B-E)** Representative images of live S2 cells that were co-transfected with Venus-Ph (B, C) or Venus-mini-Ph (D, E). H2Av-RFP was co-transfected as a nuclear marker. Images are from a single slice from a confocal stack and are arranged (1-8) based on the mean nuclear Venus intensity. Note that images were adjusted to make the signals visible for presentation, so the intensities cannot be compared across images. Scale bar is 3 microns. (C, E) Line scans through condensates to assess co-localization with chromatin. Both signals are scaled to their maximum intensity on the y-axis. (F) Relationship between mean nuclear Venus intensity and condensate formation. Cells without condensates are indicated in magenta, and those with condensates in black. Bar shows median. WT-Ph, n=391; mini-Ph, n=93.

In vitro, mini-Ph forms condensates through phase separation, a process with a sharp concentration dependence. To assess the relationship between condensate formation and protein concentration, we plotted mean intensities of at least 48 nuclei, and coded them based on whether they had at least one condensate (**Figure 2F**). For each construct, we display 8 cells from the same image with increasing intensities (**Figure 2B, D** cells1-8). This analysis shows that Ph forms condensates even at the lowest concentrations we measured. In contrast, mini-Ph only formed condensates in cells with the highest intensities (**Figure 2D, F**). This is consistent with removing the IDRs increasing the threshold for condensate formation, although we cannot assess the threshold for Ph from our current data.

Ph is a chromatin bound protein. To determine whether condensates are likely to be chromatin associated, we plotted the profile of line scans through large condensates formed in cells with the highest expression levels for both Ph and mini-Ph (**Figure 2C, E**). This indicates that while the small condensates formed by Ph overlap chromatin, the large mini-Ph condensates exclude chromatin (also clearly visible in images 5-8 **Figure 2D**). Finally, we observed cytoplasmic condensates in ∼30% of Ph-expressing cells (e.g. cells 1, 3, 4, **Figure 2B**), while we did not observe cytoplasmic condensates in cells expressing mini-Ph. Thus, removing the Ph IDRs increases the protein concentration required for nuclear condensate formation, and leads to formation of large condensates that exclude chromatin. Mini-Ph, like Ph, does not appear to enter the nucleolus.

### Removal of IDRs affects Ph condensate size, number, and morphology

For each IDR, we compared condensates formed when the IDR is deleted, versus when it is the only IDR present (**Figure 3-5**).

**Figure 3.**
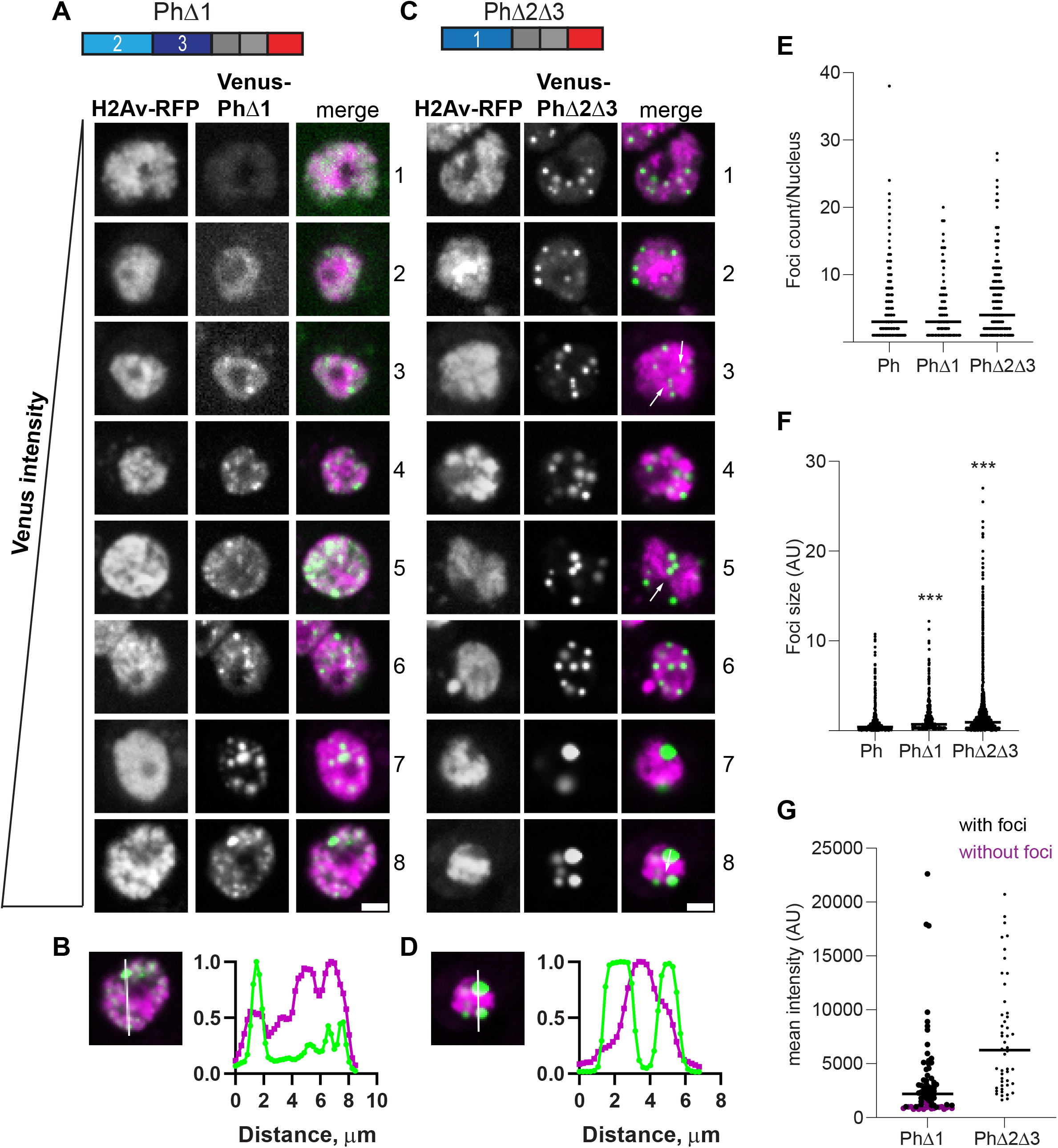
Effect of Ph1 IDR on condensates. **(A-D)** Representative images of live S2 cells that were co-transfected with Venus-PhΔ1 (A, B) or Venus-PhΔ2Δ3 (C, D). H2Av-RFP was co-transfected as a nuclear marker. Images are from a single slice from a confocal stack and are arranged based on the mean nuclear Venus intensity. Images were adjusted to make the signals visible for presentation, so the intensities cannot be compared across images. Arrow indicate chromatin “fissures”. Scale bar is 3 microns. (**B, D**) Line scans through condensates to assess co-localization with chromatin. Both signals are scaled to their maximum intensity (y-axis). (**E, F**) Graph of number of condensates (foci) per nucleus (E) and condensate size (F) from CellProfiler analysis. The total number of transfected cells analyzed (with or without foci): WT-Ph, n=3478; PhΔ1, n=2426; PhΔ2Δ3, n=2911. p-values are for comparison with WT using a Kruskal-Wallis test with Dunnett’s correction for multiple comparisons. (**G**) Relationship between mean nuclear intensity and condensate formation. Cells without condensates are indicated in magenta, and those with condensates in black. Bar shows median. PhΔ1, n=78; PhΔ2Δ3, n=48.

**Figure 4.**
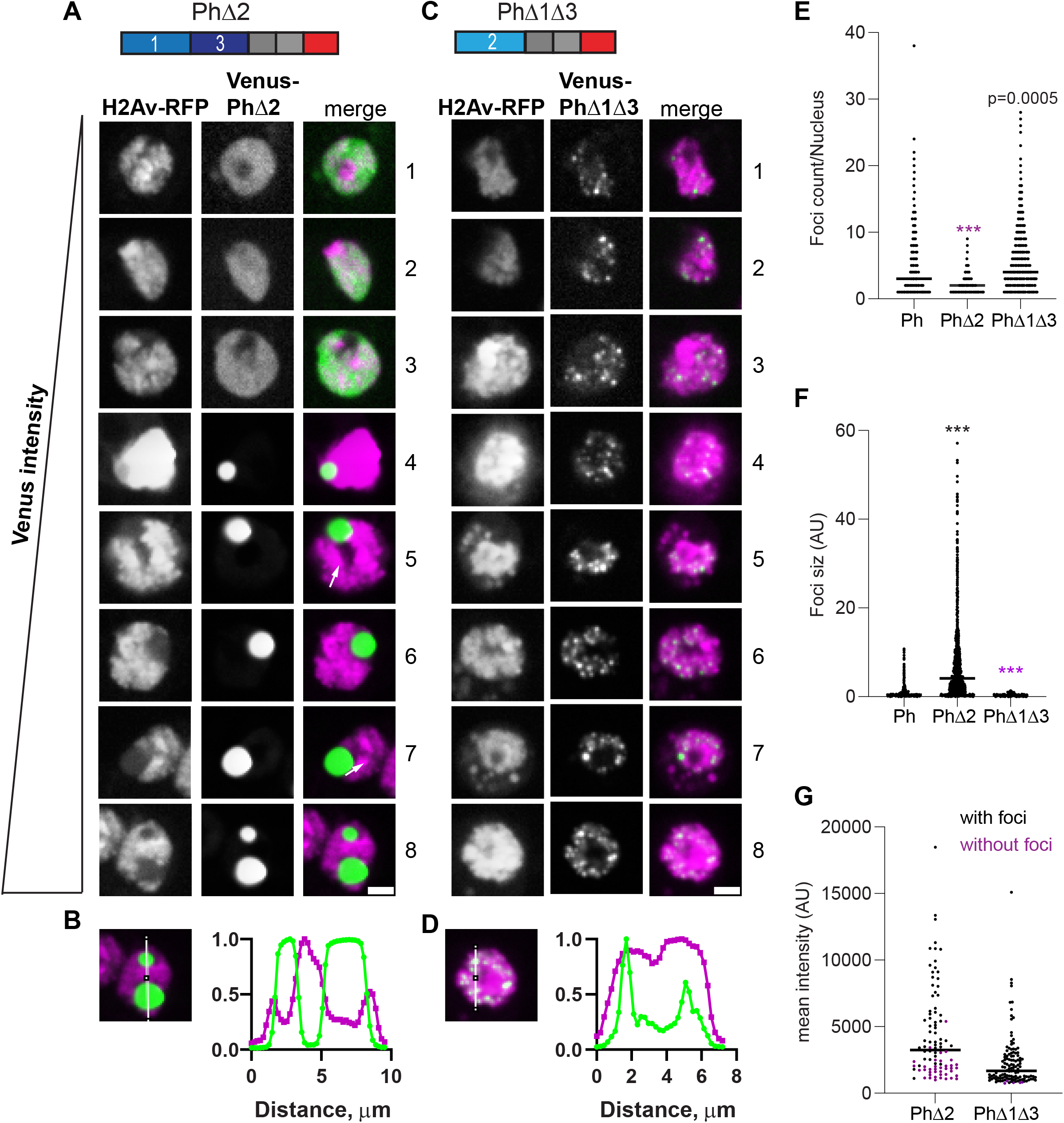
Effect of Ph2 IDR on condensates. **(A-D)** Representative images of live S2 cells that were co-transfected with Venus-PhΔ2 (B) or Venus-PhΔ1Δ3. H2Av-RFP was co-transfected as a nuclear marker. Images are from a single slice from a confocal stack and are arranged based on the mean nuclear Venus intensity. Images were adjusted to make the signals visible for presentation, so the intensities cannot be compared across images. White arrows indicate examples of chromatin “fissures”. Scale bar is 3 microns. (**B, D**) Line scans through condensates to assess co-localization with chromatin. Both signals are scaled to their maximum intensity (y-axis). (**E, F**) Graph of number of condensates (foci) per nucleus (E) and condensate size (F) from CellProfiler analysis. The total number of transfected cells analyzed (with or without foci): WT-Ph, n=3478; PhΔ2, n=1539; PhΔ1Δ3, n=1568. p-values are for comparison with WT using a Kruskal-Wallis test with Dunnett’s correction for multiple comparisons. (**G**) Relationship between mean intensity and condensate formation. Cells without condensates are indicated in magenta, and those with condensates in black. Bar shows median. PhΔ2, n=95; PhΔ1Δ3, n=124.

**Figure 5.**
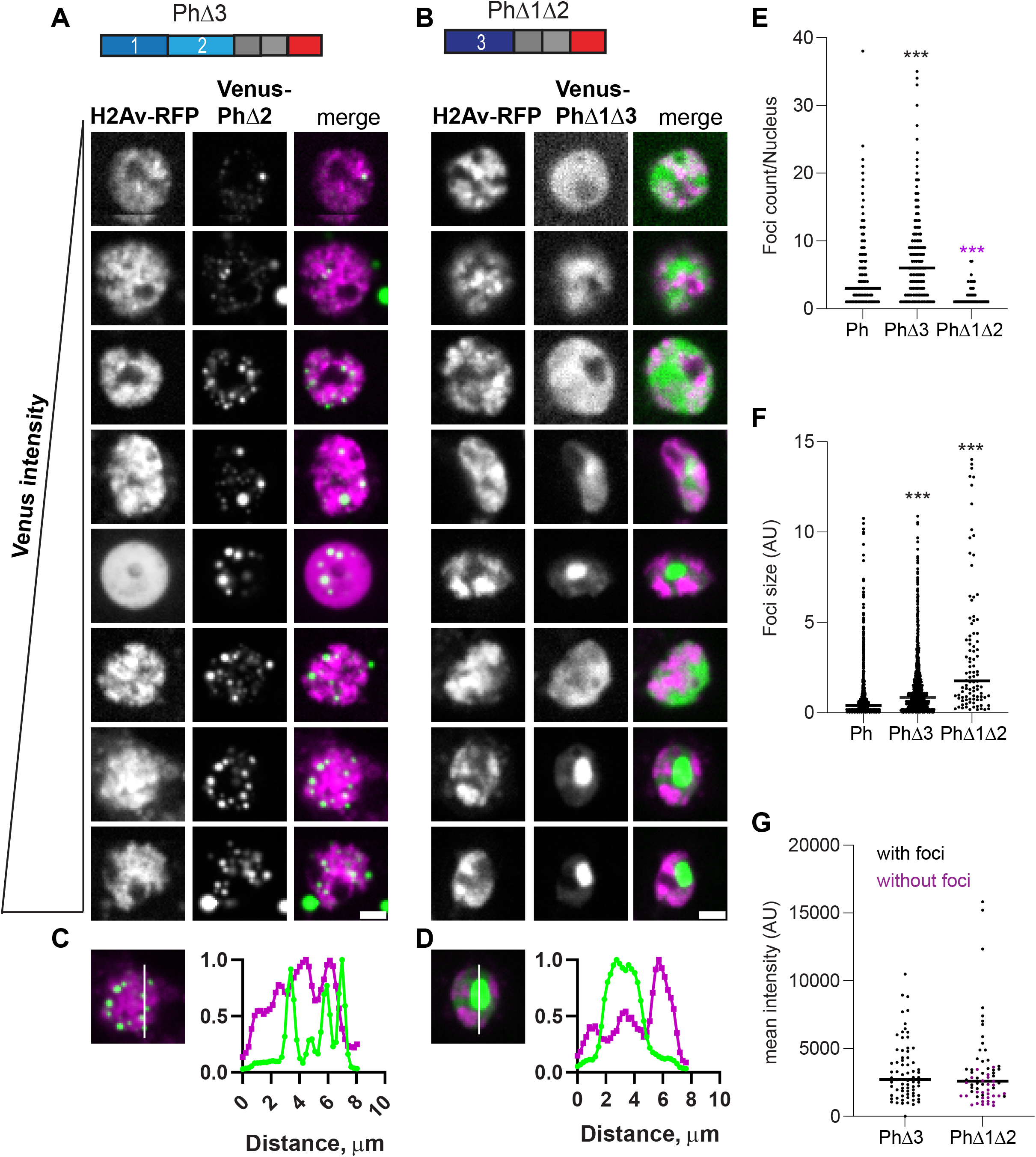
Effect of Ph3 IDR on condensates. **(A-D)** Representative images of live S2 cells that were co-transfected with Venus-PhΔ3 (B) or Venus-PhΔ1Δ2. H2Av-RFP was co-transfected as a nuclear marker. Images are from a single slice from a confocal stack and are arranged based on the mean nuclear Venus intensity. Images were adjusted to make the signals visible for presentation, so the intensities cannot be compared across images. Scale bar is 3 microns. (**B, D**) Line scans through condensates to assess co-localization with chromatin. Both signals are scaled to their maximum intensity (y-axis). (**E, F**) Graph of number of condensates (foci) per nucleus (E) and condensate size (F) from CellProfiler analysis. The total number of transfected cells analyzed (with or without foci): WT-Ph, n=3478; PhΔ3, n=1557; PhΔ1Δ2, n=1038. p-values are for comparison with WT using a Kruskal-Wallis test with Dunnett’s correction for multiple comparisons. (**G**) Relationship between mean intensity and condensate formation. Cells without condensates are indicated in magenta, and those with condensates in black. Bar shows median. PhΔ3, n=71; PhΔ1Δ2, n=67.

Condensates formed in the absence of Ph1 (PhΔ1) are heterogeneous, frequently small, and tend to form interconnected networks (**Figure 3A**). This likely explains the significant increase in foci size, but not number observed for PhΔ1 (**Figure 3E, F**). As PhΔ1 concentrations increase, condensates are more apparent, but substantial fluorescence outside condensates remains in most cells. The pattern of PhΔ1 outside of condensates appears granular, raising the possibility that PhΔ1 forms tiny clusters that cannot be resolved with our imaging method. PhΔ1 condensates overlap with chromatin (**Figure 3B**). Most cells with detectable PhΔ1 expression formed at least one visible condensate in our manual analysis, even when nuclear intensity is low (**Figure 3G**).

Condensates formed by PhΔ2Δ3 (only Ph1 present) are completely different. Round condensates are easily visible at even the lowest protein concentrations (**Figure 3C, G**). At high protein concentrations, cells with very large condensates, reminiscent of mini-Ph expressing cells, are observed (**Figure 3C**, cells 7, 8). For PhΔ2Δ3, the large condensates formed in cells with high protein levels clearly exclude chromatin (**Figure 3D**), and many smaller condensates appear to do so as well (e.g. cell 4, **Figure 3C**). We also noticed apparent chromatin “fissures” in some nuclei expressing PhΔ2Δ3 (arrows in **Figure 3C**). Condensates seem to form or collect in these fissures. PhΔ2Δ3 cells formed significantly larger foci than WT-Ph cells (**Figure 3F**).

Expression of Ph lacking Ph2 (PhΔ2), the glutamine rich IDR, results in a small number of large condensates (**Figure 4A**). Consistent with this impression, the number of foci per nucleus counted for PhΔ2 is reduced compared to wild type and there is a significant increase in foci size (**Figure 4E, F**). These condensates are similar to those formed by mini-Ph, including that they clearly excluding chromatin (**Figure 4B**). However, cells with more than one mini-Ph condensate are very rare, while cells with two condensates are common for PhΔ2. PhΔ2 expressing cells also clearly have chromatin fissures (e.g. cells 5, 7 **Figure 4A**). Like mini-Ph, PhΔ2 has a high threshold for condensate formation, so that many cells with clear Venus-PhΔ2 expression are observed without condensates (**Figure 4A, G**).

Condensates formed with Ph containing only the Ph2 IDR (PhΔ1Δ3, **Figure 4C**) are completely distinct from PhΔ2 condensates. They are significantly smaller and more numerous than WT-Ph condensates (**Figure 4E, F**), and appear embedded in chromatin (**Figure 4D**). Condensates form at even the lowest expression levels analyzed, similar to WT-Ph (**Figure 4G**).

Condensates formed in the absence of Ph3 (PhΔ3) are bright and numerous, and form at the lowest expression levels analyzed (**Figure 5**). Like WT-Ph condensates, cytoplasmic condensates are frequently present in cells expression PhΔ3 (37% cells; e.g. **Figure 5A**, cells 2, 8). PhΔ3 condensates appear most similar to WT-Ph condensates, although they are more numerous and slightly larger (**Figure 5E, F**). Nuclear PhΔ3 condensates overlap chromatin (**Figure 5B**), and form at the lowest concentrations measured (**Figure 5G**).

When Ph3 is the only IDR present (PhΔ1Δ2), the pattern of condensates is similar to mini-Ph in that cells with high protein concentrations tend to form a single large condensate that excludes chromatin (**Figure 5C, D**). Cells expressing PhΔ1Δ2, even without condensates, have a heterogeneous chromatin distribution that appears non-overlapping with PhΔ1Δ2 (**Figure 5C**). This pattern is not observed for mini-Ph or any of the other proteins analyzed. PhΔ1Δ2 has a higher expression threshold for condensate formation, like mini-Ph (**Figure 5G**).

Taken together, the data indicate that each IDR affects condensate formation. The most obvious pattern is that proteins lacking Ph2 (PhΔ2, PhΔ2Δ3, PhΔ1Δ2) consistently from large round condensates that exclude chromatin, while those containing Ph2 form small chromatin associated condensates. Thus, Ph2 has the most dramatic effect on condensate formation. Comparison of single versus double IDR deletions also indicates that the IDRs can influence each other. Specifically, Ph3 inhibits the effects of both Ph1 and Ph2. Ph1 promotes formation of small round condensates, but only in the absence of Ph3 (compare PhΔ2Δ3, **Figure 3C** to PhΔ2, **Figure 4A**). Ph2 drives formation of small condensates at low concentrations, but this is more evident in PhΔ1Δ3 than in PhΔ1 (compare **Figure 3A** to **Figure 4C**). **Figure S4** compiles results from all proteins to facilitate comparisons among the effects of the IDRs.

### Ph IDRs regulate chromatin association

To understand how the Ph IDRs affect chromatin association, we used subcellular fractionation. Transfected cells expressing Venus tagged Ph proteins cells were fractionated into cytoplasmic, soluble nuclear, and chromatin associated fractions (**Figure 6, Figure S5**). As expected, and consistent with previous (and current) analysis of endogenous Ph [46] (**Figure 6B**), Venus-Ph is mainly present in the chromatin and soluble nuclear fractions (**Figure 6C, D**). Removal of the SAM, which is required for condensate formation by full length Ph, increases the fraction of protein in the soluble nuclear fraction (**Figure 6D**). Analysis of Ph lacking IDRs indicates that proteins lacking Ph2 (mini-Ph, PhΔ2, PhΔ1Δ2, PhΔ2Δ3) are mainly present in the cytoplasmic (mini-Ph, PhΔ2), and soluble nuclear (PhΔ1Δ1, PhΔ2Δ3) fractions, with reduced levels on chromatin (**Figure 6E**). This is consistent with observed chromatin exclusion in the microscopy experiments, although we do not clearly observe cytoplasmic protein by microscopy. Proteins containing Ph2 but lacking one or two IDRs (PhΔ1, PhΔ3, PhΔ1Δ3) are more strongly chromatin associated than Ph, consistent with the apparent overlap of these condensates with chromatin (**Figure 6D**). Removal of Ph1, Ph3, or both increase the fraction of chromatin bound Ph. This is consistent with Ph3 and Ph1inhibiting chromatin association mediated by Ph2. Fractionation results are summarized in **Figure 6F**, which clearly shows how Ph2 alters the distribution of Ph in the cell.

**Figure 6.**
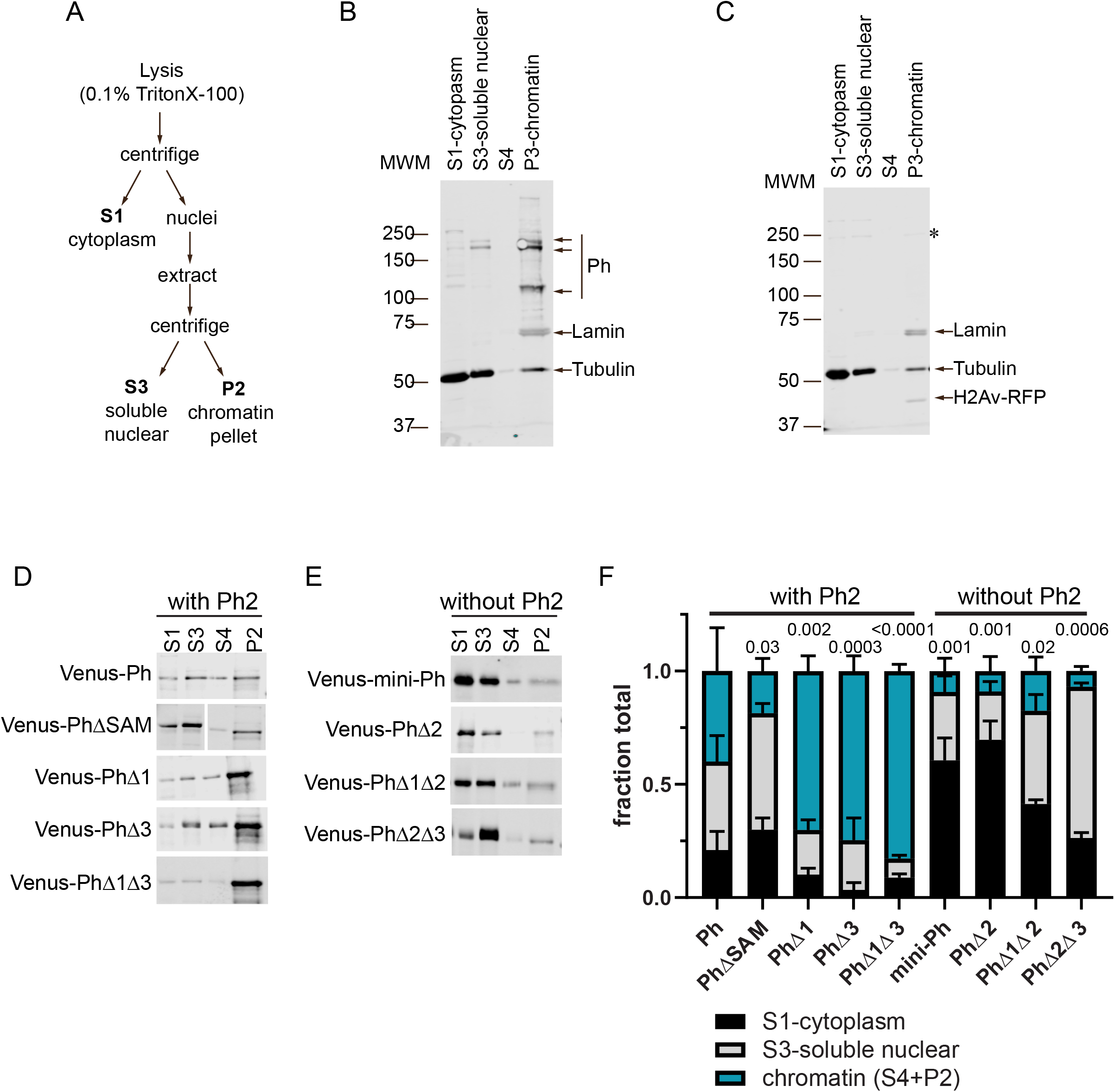
Effect of Ph IDRs on chromatin association. **(A)** Schematic of cell fractionation protocol (based on **[53]**). Note that P2 was digested with DNaseI and RNaseA, and the supernatant from this analyzed as S4 on blots. However, this digest was not successful, as histones were not released into the supernatant so that we pooled the signals from S4 and P2 for quantification. **(B)** Representative Western blot of subcellular fractionation of untransfected S2 cells. Lamin was used as a nuclear marker and Tubulin as a cytoplasmic one. Endogenous Ph (Ph-p and Ph-d, with major isoforms at ∼170kDa and ∼150kDa) is mainly found in the chromatin fraction. (**C**) Representative Western blot of subcellular fractionation of S2 cells transfected with Venus-Ph. Blots were probed with anti-GFP to recognize Venus-Ph (faintly visible, asteris), and reprobed with anti-Tubulin, anti-Lamin, and anti-RFP to detect co-transfected H2Av-RFP. (D, E) Representative Western blots of fractions of cells transfected with the indicated constructs, separated by whether they include (D) or do not include (E) the Ph2 IDR. Venus-Ph proteins were detected with anti-GFP. Note that in all Western blots, 2X more P2 was loaded than other fractions. Fractions for cells transfected with constructs lacking Ph2 were loaded at 0.67X the amount of those expressing constructs with Ph2. (F) Summary of quantification of three independent fractionation experiments. Error bars show mean and standard deviation. Numbers are p-values for one-way Anova comparing the fraction in chromatin relative to WT-Ph.

### Removal of the Ph3 IDR allows Ph2-dependent but SAM-independent condensates to form

We showed previously that the Ph SAM is necessary for condensates to form in cells [26]. To determine if the Ph IDR deletion constructs form condensates in the absence of SAM, we tested all the constructs except PhΔ1Δ2 without SAM (**Figure 7A**). In the absence of the SAM, most proteins did not form condensates. However, in two cases, deletion of Ph3 and deletions of both Ph1 and Ph3, condensates form without the SAM although the number of cells that form condensates is low (**Figure 7**). Although the ΔSAM versions are expressed at about 2-fold lower levels than the corresponding proteins with the SAM (**Figure S3**), given the wide range of expression levels over which condensates are observed, this is unlikely to explain why most ΔSAM proteins do not form condensates. The ΔSAM versions are also expressed at similar levels as transfected WT-Ph (**Figure S3**).

**Figure 7.**
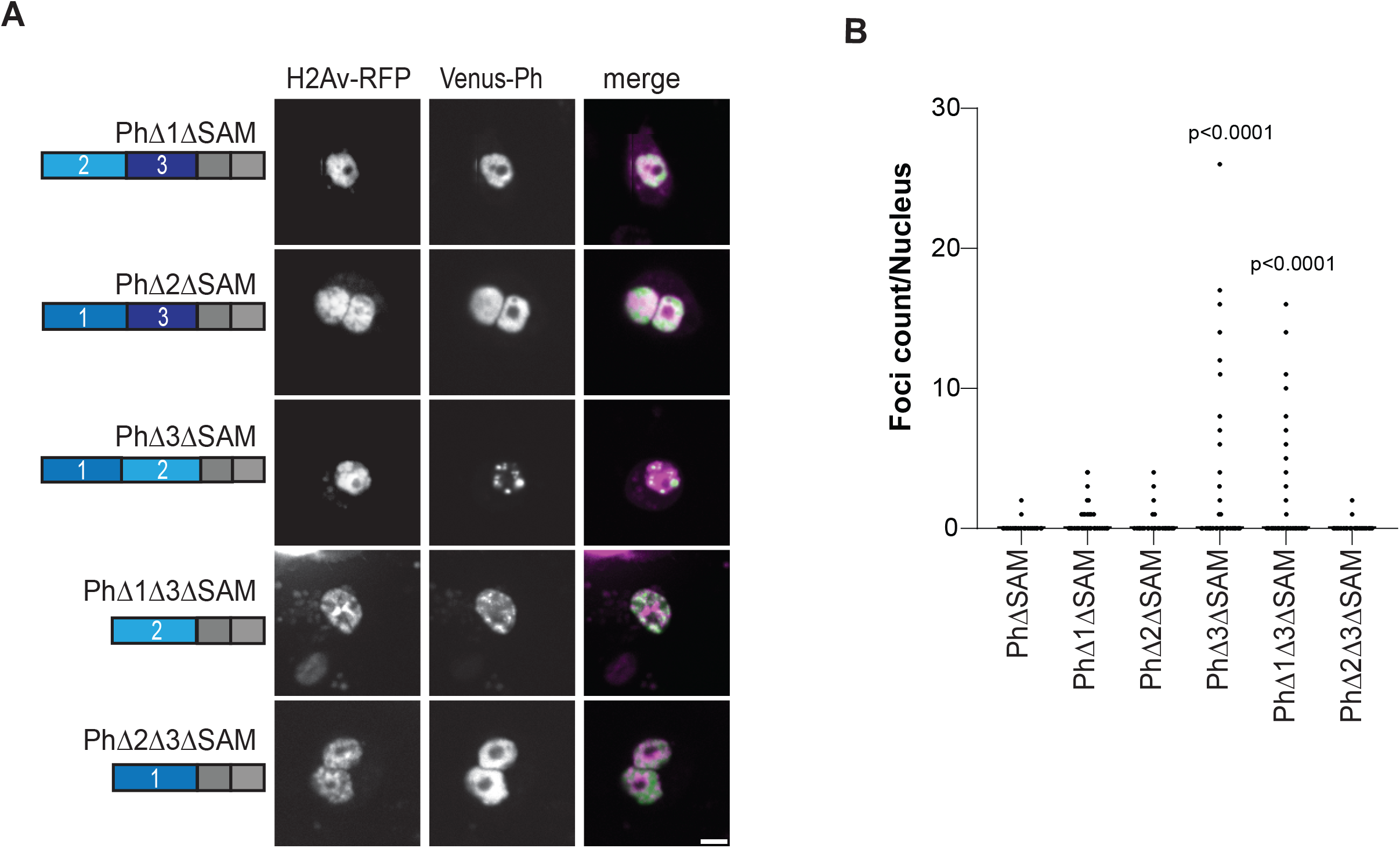
Effect of Ph IDRs on condensate formation in the absence of the SAM. **(A)** Representative live images of S2 cells transfected with Venus-Ph constructs lacking one or two IDRs and the SAM with H2Av-RFP as a nuclear marker. Images show maximum intensity projections of confocal stacks. Scale bar is 5 microns. **(B)** Quantification of foci count per nucleus for H2Av-RFP positive cells. At least 100 cells from each of the three independent experiments were analyzed. The total number of cells analyzed was: PhΔSAM, n=1348; PhΔ1ΔSAM, n=411; PhΔ2ΔSAM, n=359; PhΔ3ΔSAM, n=214; PhΔ1Δ3ΔSAM, n=922; PhΔ2Δ3ΔSAM, n=1413. p-values are for comparison with PhΔSAM using a Kruskal-Wallis test with Dunnett’s correction for multiple comparisons.

### Ph IDRs alone and in combination can form condensates

To test if the IDRs can form condensates in the absence of mini-Ph, we tested each IDR alone or in combination (**Figure 8**). An SV40 NLS was added to ensure that the IDRs localize to the nucleus, since the Ph NLS is in the FCS domain. The proteins were expressed within 2X the level of transfected WT-Ph, and total Ph levels were not increased more than 2X (**Figure S6**). Ph3 alone does not form condensates. However, both Ph 1 and Ph2 form condensates in a small number of cells (**Figure 8B**). When Ph1 and Ph2 are combined (Ph5), many small condensates are formed (**Figure 8B**). In contrast, when Ph2 and Ph3 are combined (Ph6), condensates do not form. Finally, when Ph1, Ph2, and Ph3 are combined (Ph7), condensates are formed, although fewer than with Ph1+Ph2 (Ph5) (**Figure 8B**). These results indicate that intrinsic activity of the IDRs may contribute to Ph condensates, but may also be constrained by the mini-Ph region. The finding that Ph5 frequently forms condensates, while Ph6 doesn’t and Ph7 rarely does, confirms the inhibitory effect of Ph3 on Ph1 and Ph2. **Table 1** shows a comprehensive summary of condensate formation for all of the constructs tested.

**Figure 8.**
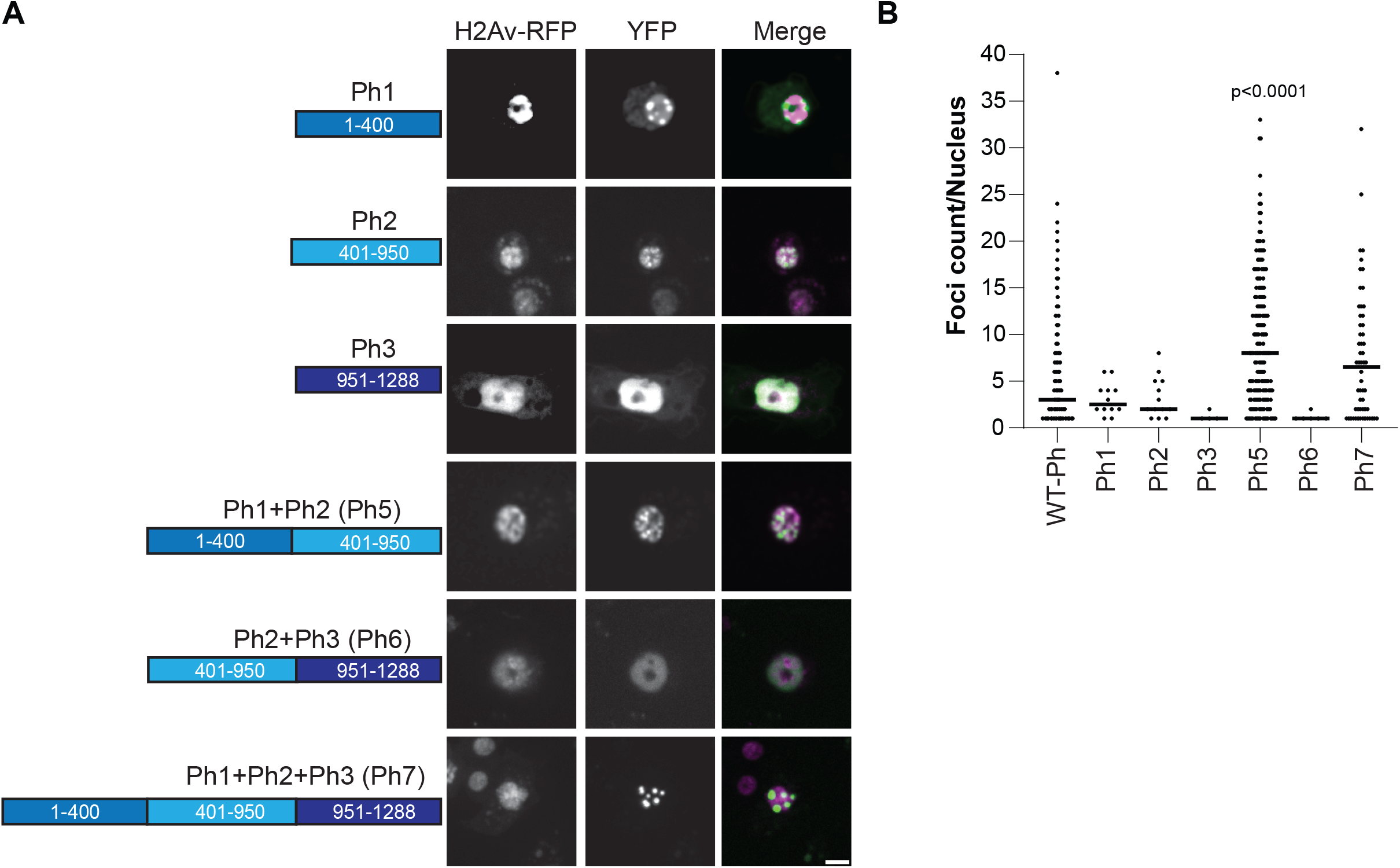
Condensate formation by Ph IDRs. **(A)** Representative live images of S2 cells transfected with the Venus-Ph truncation constructs (IDRs only) with H2Av-RFP as a nuclear marker. Images show maximum intensity projections of confocal stacks and scale bar is 5 microns. **(B)** Quantification of foci per nucleus. Bars show the median value for cells that formed foci, pooled from three independent experiments. The total number of transfected cells analyzed (with and without foci) were as follows: Ph1, n=268; Ph2, n=412; Ph3, n=309; Ph5, n=857; Ph6, n=460; Ph7, n=175. p-values are for comparison with WT-Ph using the Kruskal-Wallis test with Dunnett’s correction for multiple comparisons.

**Table 1.** Summary of condensate formation by Ph proteins.

| CONSTRUCT | SAM | CONDENSATES |
| --- | --- | --- |
| PhΔ1 | + | + |
| PhΔ2 | + | + |
| PhΔ3 | + | + |
| PhΔ1Δ3 | + | + |
| PhΔ2Δ3 | + | + |
| PhΔ1Δ2 | + | - |
| PhΔ1ΔSAM | - | - |
| PhΔ2ΔSAM | - | - |
| PhΔ3ΔSAM | - | + |
| PhΔ1Δ3ΔSAM | - | + |

In summary, we have identified three IDRs in the N-terminal region of Ph, each of which influence Ph SAM-dependent condensates in cells. The most dramatic effects occur when the glutamine rich Ph2 IDR is removed, which reduces condensate binding and leads to formation of large, round condensates. The Ph3 IDR, which was previously shown to inhibit aggregation of the SAM through its o-linked glycosylation, also inhibits Ph1 and Ph2 activities. The effects of the IDRs correlate with their effects on chromatin association—weak chromatin association corresponds to large, round condensates, while strong chromatin association correlates with small, chromatin associated condensates. **Figure 9** summarizes these results.

**Figure 9.**
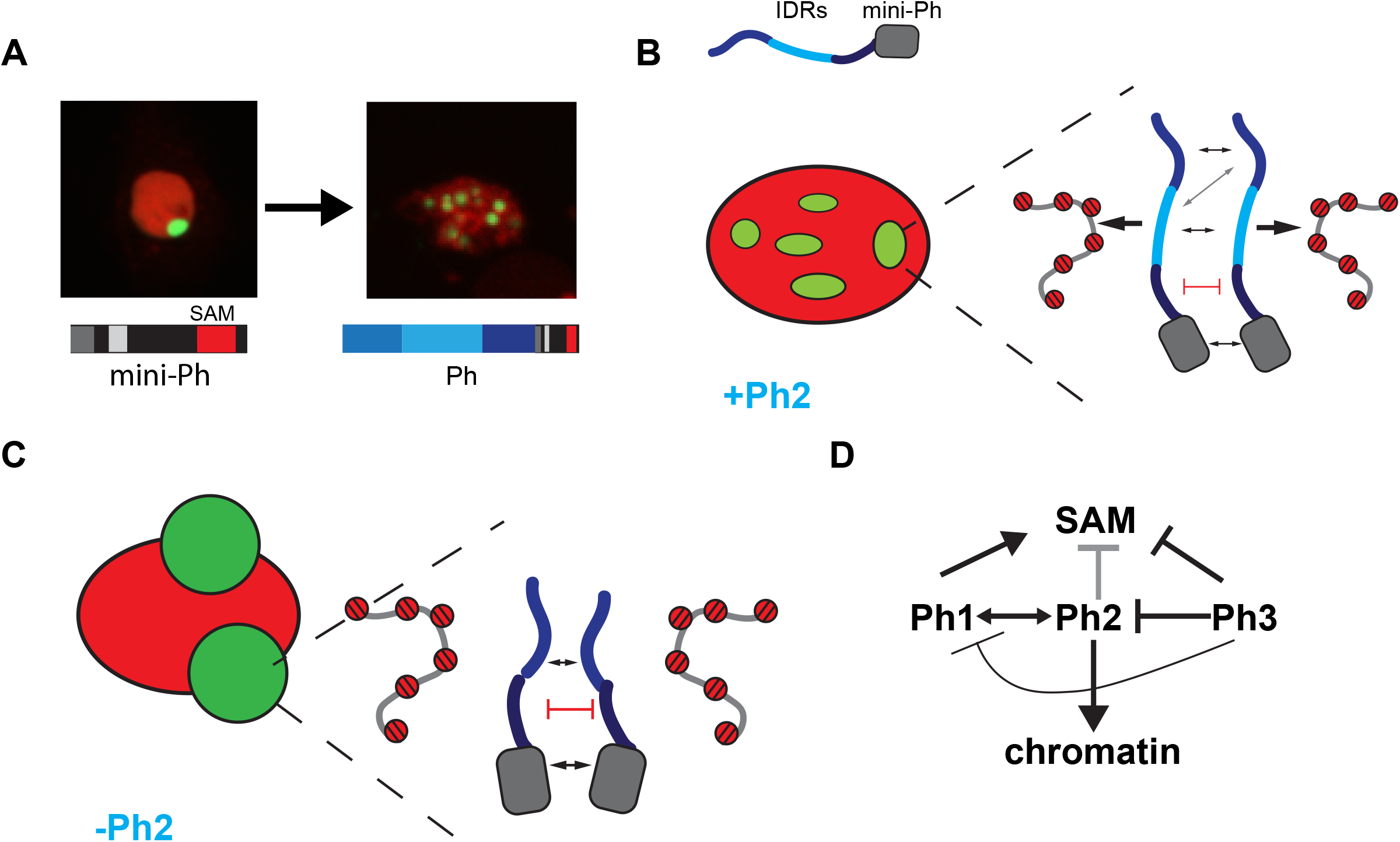
Model for role of Ph IDRs in SAM dependent condensate formation. **(A)** Comparison of condensates formed by mini-Ph and Ph demonstrates the effect of the IDRs. (**B**) Schematic of interactions that may be balanced to form multiple condensates in constructs containing the Ph2 IDR. Protein-protein interactions between the mini-Ph region, and among the IDRs are predicted to drive phase separation, while tight binding to chromatin mediated by Ph2 may restrict condensate formation. (**C**) Schematic of how removing Ph2 may lead to formation of large condensates by removing the constraint imposed by tight chromatin binding. (D) Working model for regulatory network among IDRs. Ph2 (gray arrow) restricts both the size of condensates and propensity for formation when Ph SAM is present, and imparts tight chromatin association. Ph3 inhibits the ability of both Ph1 and Ph2 to promote condensate formation; it may also inhibit the SAM (as proposed previously).

## DISCUSSION:L

We sought to determine how SAM dependent condensates that may form by phase separation are controlled by the disordered N-terminal region of Ph. We identified three distinct IDRs in this region by sequence analysis and tested the function of each, in the context of full length Ph, Ph without the SAM, and alone. We find that each IDR affects condensates and binding to chromatin, and that the IDRs interact functionally with each other. The potential mechanisms underlying these effects are considered below.

Ph1 promotes condensate formation. When Ph1 is fused to mini-Ph (PhΔ2Δ3), small round condensates are formed that are wild-type like (although they are more numerous than WT-Ph) (Figure 3). These condensates are dependent on the Ph SAM (Figure 7). When Ph1 is deleted, condensates are less round, and diffuse granular staining is often observed. Ph1 alone can form condensates in cells, although it does so rarely (Figure 8). Taken together, these data suggest Ph1 promotes the condensate forming ability of Ph SAM, leading to formation of many condensates in a high fraction of cells.

Ph2 has the most profound effects on condensates. In the presence of Ph SAM, it controls condensate size, so that its removal leads to formation of very large condensates, similar to those formed by mini-Ph (Figure 4). Ph2 fused to mini-Ph results in many small condensates (although in a small number of cells). Ph2 promotes condensate formation in the absence of the SAM, but only if Ph3 is absent (Figure 7). Ph2 alone has weak condensate forming ability, but when fused to Ph1 forms a large number of small condensates (Figure 8). Thus, Ph2 negatively regulates Ph SAM dependent condensates, but also interacts functionally with the other IDRs in a SAM independent manner—it synergizes with Ph1 to promote condensates, and is inhibited by Ph3. The most striking sequence feature of Ph2 is its high glutamine (36%) content, and presence of prion-domains (Figure 1). It is also intriguing to note that a frequently used *ph* allele, *ph*^*505*^, which was thought to be a null, actually encodes truncated versions of both *ph-p* and *ph-d*. The predicted proteins are truncated midway through the Ph2 region, so would potentially produce proteins similar to Ph5 (i.e. Ph 1+2). *ph*^*505*^ effects on gene regulation differ slightly from those of a true *ph* null **[47]**, raising the possibility that *ph*^*505*^ effects are due to expression of the Ph5-like protein. Although we refer to Ph2 as an IDR, its glutamine rich regions are predicted to have helix forming potential. Thus, future studies should assess a possible contribution of helix-formation in the activity of Ph2, particularly in light of recent work implicating helix-forming potential of glutamine rich sequences in regulating phase separation [37]. Ph3 has an inhibitory effect on condensates so that more condensates are formed when it is removed. Careful consideration of the effect of removing and having Ph3 in different constructs suggests Ph3 may function through the other IDRs, particularly by restricting the activity of Ph2. First, when Ph3 is fused to mini-Ph (PhΔ1Δ2), few condensates are formed, and the ones that form are quite similar to mini-Ph alone, suggesting Ph3 has little effect in this context (although it appears to globally affect chromatin organization) (Figure 5). Second, Ph1 fused to mini-Ph (PhΔ2Δ3) forms several condensates (Figure 3), but the number is significantly reduced (and size increased) when Ph3 is present with Ph1 (PhΔ2) (Figure 4). Third, Ph2 forms condensates in the absence of SAM (PhΔ1Δ3ΔSAM) but with Ph3 present does not (PhΔ1ΔSAM) (Figure 7). Fourth, Ph2 alone forms condensates in some cells but again condensate formation is inhibited when Ph2 is fused to Ph3 (Ph6) (Figure 8). Thus, Ph3 function may be through the other IDRs, rather than directly on Ph SAM; alternatively, it could inhibit both Ph SAM and Ph2. Ph3 is rich in serine and threonine and is known to be extensively modified by O-linked glycosylation of these residues [31] as well as phosphorylation **[48]**. Removing glycosylation drives formation of Ph aggregates *in vitro* and *in vivo*, and these aggregates depend on the SAM [25]. Whether these aggregates also depend on Ph2 is not known. However, one hypothesis would be that glycosylation restrains the activity of Ph2 that promotes condensate formation through SAM-dependent and independent mechanisms. Recently, O-GlcNacylation was also shown to reduce both aggregation and phase separation of the N-terminal LCR of the EWS protein **[49]**; future experiments will test if this is also true for Ph. It should be noted that a protein very similar to PhΔ3 was expressed in *Drosophila* imaginal discs, and formed condensates similar to the wild-type protein, consistent with our observations in S2 cells [25].

Many mechanisms have been described and hypothesized to explain control of condensate size and number in cells **[50–52]**. One model that may be relevant for Ph1 and Ph2 explains how chromatin binding affects condensate number and size. Qi and Zhang used simulations to show that protein-chromatin interactions can promote a multi-droplet state **[52]**. In this model, protein-chromatin contacts promote nucleation of phase separation, but inhibit their coalescence. Detailed investigation of the thermodynamics underlying these effects indicate that this is due to a kinetic barrier between single and multi-droplet states. This kinetic barrier arises from the chromatin network—when droplets that interact with chromatin coalesce, the chromatin network is constrained, which becomes progressively more energetically costly. Theoretical work from Wingreen and colleagues also finds that chromatin interferes with coarsening of condensates, both by acting as a crowder, and through its crosslinked network **[51]**. Thus, changing protein-chromatin interactions could change droplet number. In the framework of these models, Ph1 may decrease chromatin binding affinity, promoting formation of many round condensates. Loss of Ph1 may lead to tight chromatin binding, explaining the distorted shape of condensates formed by PhΔ1, and consistent with our chromatin fractionation data. Ph2, on the other hand, promotes chromatin binding so that its loss leads to large condensates because fusion is not impaired.

Shin et al. **[53]** analyzed condensate formation by chromatin proteins using optogenetic manipulation to induce clustering/phase separation. They find that condensates of different chromatin proteins, as well as endogenous condensate forming nuclear proteins, occupy regions of low chromatin density. This is consistent with what we observe with large Ph condensates.

Shin et al. **[53]** also showed that specific chromatin regions that interact with the phase separating protein associate with condensates, leading to a model where condensates can act as a filter to bring specific chromatin regions together while excluding bulk chromatin **[53]**. Thus, an important next step will be to determine how the Ph IDRs affect Ph binding to specific PcG targets in chromatin.

Recently, Seydoux and colleagues have demonstrated that clusters of MEG-3 function as Pickering particles, which adsorb to the surface of P-granules and reduce their coarsening by reducing surface tension **[54]**. As speculated, RNA may also have this function for some condensates **[54]**. In this regard it is interesting that the Ph2 region is predicted (using DisoRDPbind **[55]**) to have a strong RNA binding activity, although this has not been validated experimentally.

A caveat to this work is that we studied behaviour of Ph proteins under conditions of overexpression, and with endogenous Ph present in the background. While this allows us to evaluate how the different IDRs affect Ph condensate formation in a cellular context, we do not know which effects are dependent on endogenous Ph. Total Ph levels from transfected cells were not more than 2.5X of untransfected cells, due to downregulation of endogenous Ph (Figure S3E, F). All of the constructs except the Ph IDRs alone (Figure 8) should assemble into PRC1, since this is mediated by the HD1 domain. However, we cannot rule out that some IDR effects are due to interactions with cellular proteins. These interactions may be saturated in cells with high expression of the ectopic Ph proteins, but overexpression can also lead to spurious interactions. Our experiments reveal the complex effects of the IDRs; determining how these properties affect Ph and its function in gene regulation under normal physiological conditions awaits additional experiments. This cell-based characterization of the Ph IDRs also motivates future *in vitro* and *in silico* analysis to understand which sequence properties drive the effects of the IDRs, and how activities like DNA (or RNA) binding and phase separation contribute. We showed previously that mini-Ph undergoes phase separation in vitro; some of the condensates observed here are round and can be observed to fuse, properties consistent with (but not diagnostic of) phase separation. However, whether the small condensates driven by the presence of Ph2 form through a different mechanism awaits further study.

In conclusion, we showed that the disordered N-terminal region of Ph, which was largely uncharacterized, affects Ph SAM-dependent condensate formation, likely by affecting interaction with chromatin. The complex effects of the IDRs on condensates, and the functional interactions among them, indicate that condensate formation by the PcG is a tightly regulated process that integrates IDRs and structured regions like the SAM. Most isoforms of all three mammalian Ph homologues have disordered N-terminal IDRs; we hypothesize that IDR regulation of condensate formation by these IDRs may be functionally conserved.

## MATERIALS AND METHODS

### Cloning

Ph, and all of the truncations and deletions were cloned into a gateway donor vector using restriction digest and ligation, PCR, or gene blocks spanning deletion junctions. No extra sequence was added—for example, PhD2 consists of aa1-400-951-1589. All synthetic and PCR generated sequence was confirmed by Sanger sequencing. The plasmid for Ph truncations (pCR8-ATG-NLS) includes an SV40 nuclear localization sequence (NLS). To transfer Ph truncations into plasmid pHVW (DGRC stock # 1089) for heat-shock inducible expression in *Drosophila* cells, LR-recombination reactions were performed with 75ng of donor and 75ng of acceptor in a 5μl reaction, according to the manufacturer’s protocol. Plasmids for transfection were prepared by Qiagen maxiprep.

### Cell Culture

*Drosophila* Schneider 2 (S2) cells (Expression Systems, 94-005F) were cultured in ESF media (ESF 921 Insect Cell Culture Medium, Expression Systems) with 5% fetal bovine serum (FBS, Wisent) at room temperature on plates. Cells were passaged every 2 to 3 days. night before transfection, 1.5e6 S2 cells were plated per well of a 6-well plate. The next day, the media was changed, and transfection mix was added. Mirus Transit insect Transfection Reagent (Mirus bio) was used for the transfections according to the manufacturer’s protocol. To mark nuclei, pAct5C-H2Av-RFP (gift of V. Archambault) was co-transfected with Venus constructs. The day after transfection, the media was changed, and the next day, cells were replated on a ConA-coated glass-bottom imaging dish (Ibidi). Cells were heat shocked for 8 min at 37°C to induce Ph expression, and used the next day for live imaging.

### Western Blotting

500,000 S2 cells per well were plated in a 24 well plate (Corning 353047) one day prior to transfection. Fresh media was added the next day and transfection was carried out as above. The day after, the media was changed and the evening of next day, cells were heat shocked for 8 min at 37°C. Post 24 hours, transfected S2 cells were counted and 500,000 cells were centrifuged at 2,500 rpm at 4°C for 5 min. Pellets were re-suspended in 70 μl 2X SDS-PAGE buffer (232μl/ml Tris pH 6.8, 100μl/ml glycerol, 34mg/ml sodium dodecyl sulfate (SDS), 120mg/ml bromophenol blue) and boiled for 5 min. Samples were then run on 8% and 16% SDS-PAGE gels for Ph and H2A-RFP respectively for 80 min at 120 Volts, and transferred to nitrocellulose membranes.

Membranes were blocked for 30 min in 5% milk/PBST (1XPBS, 0.3% TWEEN® 20) and incubated overnight at 4°C on a shaker in primary antibody diluted in 5% milk/PBST. Primary antibodies used are as follows: anti-α-tubulin (mouse, 1:3 000, Sigma Aldrich T5168), anti-Ph (rabbit, 1:3 000, Francis lab), anti-GFP (rabbit, 1:3 000, Protein tech 50430-2-AP), anti-RFP (rabbit, 1:3 000, St. Johns Laboratory STJ97083) and anti-H2B (mouse, 1:3 000, Abcam, ab52484). Membranes were washed 3 times for 10 min each in PBST, incubated for 2 hrs in secondary antibody diluted in 5% milk/PBST and washed 3 times again for 10 min in PBST. Secondary antibodies were conjugated to Alexa Fluor 680 (anti-rabbit and anti-mouse, Invitrogen A21076 and A21057 respectively) or 800 CW (anti-Rabbit, Li-Cor) and used at 1:25,000 in 5% milk/PBST. Blots were scanned on an Odyssey CLx imager.

### Cellular Fractionation

Cellular fractionation was carried out as in **[46, 56]**. Transfections were carried out as described above, but in 6-well plates. Cells were allowed to grow up to 6 days after transfection, and heat-shocked to induce expression as described above 16-20 hours before harvesting. Between 1 and 1.7e7 cells were used for each fractionation. All centrifugations were carried out at 4°C, and all procedures on ice. Cells were pelleted at 1300*g in a JS5.3 rotor for 4 min., washed with 1 ml ice cold PBS; 5% of cells were removed for total cell extracts, and the remaining cells were centrifuged as above. Pellets were resuspended in 500μl Buffer A (10mM Hepes, pH 7.9, 10mM KCl, 1.5mM MgCl_2_, 0.34M sucrose, 10% glycerol, 0.1% TritonX-100, 1mM DTT) with protease and phosphatase inhibitors as indicated (0.2mM PMSF, 10μg/ml Aprotinin, 10μg/ml Leupeptin, 2μg/ml Pepstatin, 16 μg/ml Benzamidine, 10μg/ml Phenanthroline, 50μg/ml TLCK, 50mM NaF, 2mM Na-ortho-vanadate, 40mM β-glycerophosphate). Cells were incubated on ice for 5 min., and pelleted for 4 min. at 1300*g.

Supernatant was collected as S1; pellets were washed once in 500μl Buffer A, and pelleted again. Pellets were resuspended in 500μl Buffer B (3mM EDTA, 0.2mM EGTA, 1mM DTT), with the same phosphatase and protease inhibitors as indicated for Buffer A, and incubated 30 min. on ice. Nuclei were pelleted by centrifugation at 1700*g, and supernatant collected as S3. Pellets were resuspended in 350μl DNaseI digestion buffer (10mM Tris, pH 7.6, 2.5mM MgCl_2_, 0.5mM CaCl_2_) with 20μg RNaseA and 2μl DnaseI per sample and incubated on ice for 1 hour. Nuclei were centrifuged at full speed in a refrigerated microfuge. Supernatant was saved as S4, and chromatin pellets resuspended in 2X SDS-sample buffer as P2. Analysis of histone distribution in fractions indicates that the nuclease digestion was not successful, since histones were not released into S4. We therefore pooled signals from S4 and P2 as “chromatin associated”. 6X-SDS-sample buffer was added to S1, S3, and S4 to 1X. All samples were boiled at least 5 min., and stored at -20. Western blot analysis was carried out as described above, using 10% SDS-PAGE.

### Image Acquisition

Live images were acquired on a Zeiss microscope, equipped with a Yokogawa CSU-1 spinning-disk confocal head. A 63X oil objective was used and the software for image acquisition was Zen 2012. The excitation wavelengths for Venus and RFP were 488, 561 nm, respectively. For the VENUS channel, the laser power and exposure were set as follows 2.40% and 77mS, respectively. For the RFP it was 11% and 500mS, respectively. For live imaging, 3*3 tiles of confocal stacks of 1μm slices were collected.

### Image Processing and Analysis (CellProfiler)

Live images were opened using ImageJ (Fiji) in tiff format. A maximum intensity projection was made for each image. Images were then split into VENUS and RFP channels, which were named as foci and nuclei, respectively. These were then uploaded to Cell Profiler (3.1.9). Cell Profiler modules were used to build the basic analysis pipeline but a few parameters were modified for different constructs. The basic pipeline modules were identification of nuclei from RFP staining and foci identification from the VENUS channel. The module IdentifyPrimaryObjects was used for the identification of the nucleus, which was named “nuclei”. Objects outside a diameter range and touching the image border were excluded. Global thresholding was applied with either Minimum Cross Entropy or Otsu algorithms.

Identification of foci was done using the IdentifyPrimaryObjects module with thresholding using Otsu or Minimum Cross Entropy, and named “foci”. The size of the smoothing filter and the distance between local maxima parameters were adjusted for each construct separately to segment clumped objects. In the case of constructs like PhΔ1 where extensive clumping was observed, EnhanceorSuppressFeatures was used to ignore highly clumped objects. The RelateObjectsModule was used to count the number of foci per nucleus. MeasureObjectIntensity was used to obtain mean intensity values. For the mean VENUS intensity, the nuclei objects corresponding to the nuclei images were selected. MeasureObjectSizeShape provided the measurements of size which is the total number of pixels within the object. The object selected in this case was foci. For condensates that have complex morphologies, are small, or are faint relative to surrounding levels (such as those formed by PhΔ1), the software does not detect them well, and often fuses multiple tiny condensates, so the numbers are likely underestimates.

For live image analysis, at least 100 cells were analyzed per experiment for each construct. All observations were made from at least three experiments. Statistics were calculated using GraphPad Prism v8.4.3, using recommended settings, including for correction for multiple comparisons. p values were calculated for Kruskal–Wallis tests with Dunn’s multiple-comparison correction.

### Manual Image analysis (ImageJ)

To test the relationship between protein levels and condensate formation, and to confirm results from automated analysis, we analyzed between 48 and 480 cells using a single confocal slice in ImageJ (Fiji). We selected cells with visible signal in both the Venus and H2Av-RFP channels. Nuclei were segmented by hand using the magic wand tool. In less than 10% of cases we instead used the freehand drawing tool if the magic wand could not capture the nucleus (typically if two cells are close together). The mean intensity in nuclei was then measured in the Venus channel. To count condensates, we used “find maxima”, and assigned them to each measured nucleus as described here: https://microscopy.duke.edu/guides/count-nuclear-foci-ImageJ. The find maxima “prominence” parameter were adjusted for each construct--in general, constructs that form small foci are well captured with prominence set to 1000, while 5000 or 10000 was used for constructs that form large condensates (to prevent identification of multiple maxima in single condensates). These data were used to plot mean intensities for different constructs, to compare intensity for cells with and without condensates. The same data were used to create the image galleries shown, where 8 cells from the same image were selected and ordered by mean intensity to illustrate the relationship between total protein levels and condensate formation. The images were adjusted so that both the H2A-RFP and Venus signals are visible in all cases, so the intensity cannot be compared based on the images shown. The 7th or 8th most intense cell was used to plot the profile of Venus and H2A-RFP intensity across at least one condensate (a line was drawn through the condensate and “plot profile” used in both channels), to qualitatively assess colocalization of condensates with chromatin.

## Supplementary Materials

The following supporting information is included: Figure S1 Predicted disorder in Polyhomeotic-like Ph homologues; Figure S2 Complete sequences of Ph IDRs; Figure S3 Transfected proteins are (over)expressed as full-length proteins; Figure S4 Summary of effects of Ph IDRs on condensate formation; Figure S5 Full gels of Western blots of cell fractionation; Figure S6 Quantification of Ph IDR expression levels in transfected cells.

## Author Contributions

IK prepared constructs, carried out transfections, imaging, data analysis, prepared figures and contributed to writing the manuscript. ELB prepared and validated constructs, and provided input on the manuscript. NJF supervised the project, collected and analyzed data, wrote the manuscript and provided funding. IK and NJF conceived the project.

## Data Availability Statement

The data sets generated during the current study are available from the corresponding author on reasonable request.

## Acknowledgments

IK thanks members of the Francis lab, particularly Djamouna Sihou, for critical input on the project.

