## Supplementary material for "Regulation of Polyhomeotic condensates by intrinsically disordered sequences that affect chromatin binding": 6 supplemental figures

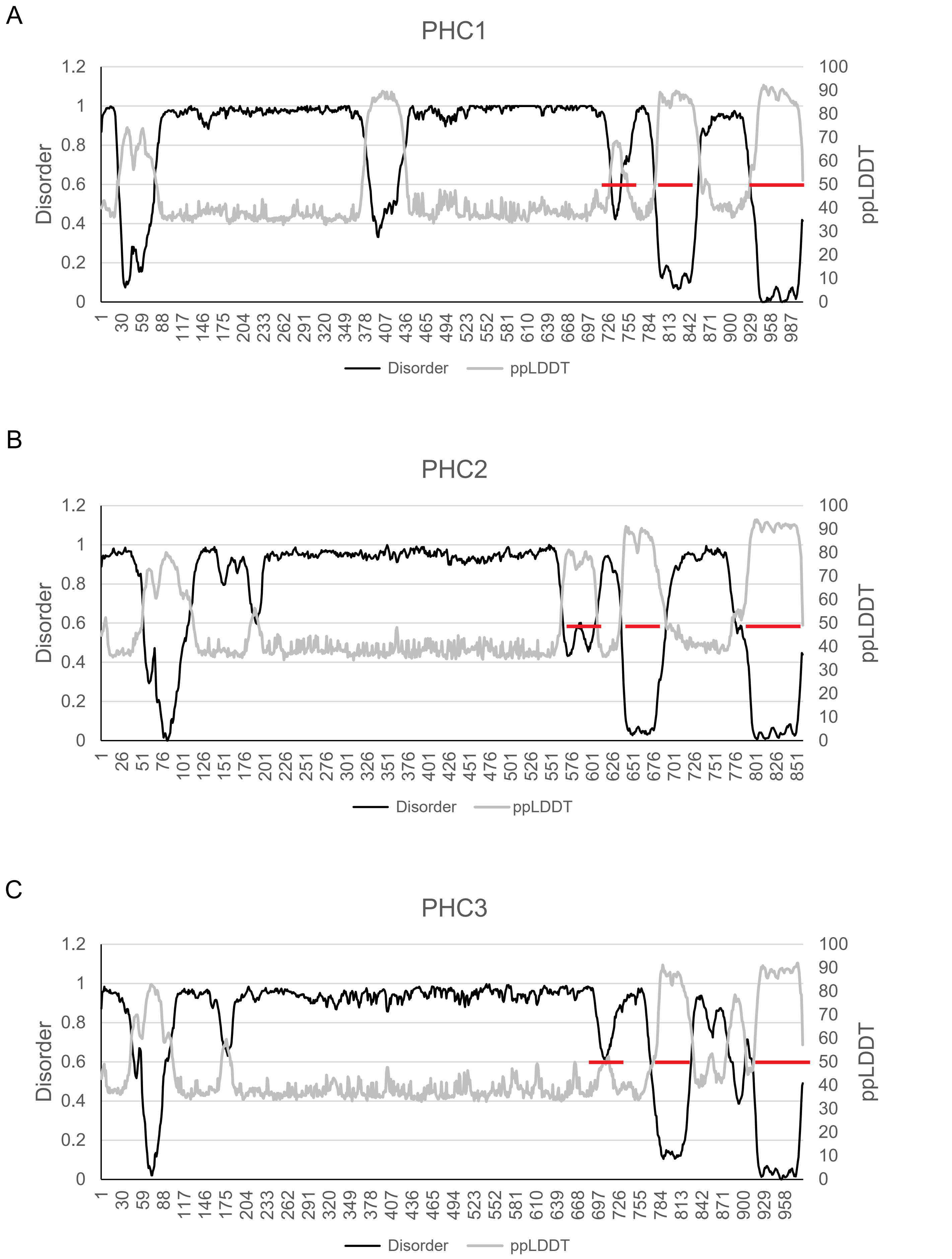


Kapur et al., Supplementary Figure 1

**Figure S1 Predicted disorder in Polyhomeotic-like Ph homologues.** (**A-C**) Metapredict2 analysis of PHC1-3. Red lines indicate positions of SAM, FCS, and HD1 domains, from C- to N-terminus. X-axis is amino acid position.

positively charged
negatively charged
polar
aliphatic
aromatic
proline

**Ph 1**

MDRRALKFMQ KRADTESDTT TPVSTTASQG ISASAILAGG TLPLKDNSNI
REKPLHHNYN HNNNNSSQHS HSHQQQQQQ VGGKQLERPL KCLETLAQKA
GITFDEKYDV ASPPHPGIAQ QQATSGTGPA TGSGSVTPTS HRHGTPPTGR
RQTHTPSTPN RPSAPSTPNT NCNSIARHTS LTLEKAQNPG QQVAATTTVP
LQISPEQLQQ FYASNPYAIQ VKQEFPTHTT SGSGTELKHA TNIMEVQQQL
QLQQQLSEAN GGGAASAGAG GAASPANSQQ SQQQQHSTAI STMSPMQLAA
ATGGVGGDWT QGRTVQLMQP STSFLYPQMI VSGNLLHPGG LGQQPIQVIT
AGKPFQGNGP QMLTTTTQNA KQMIGGQAGF AGGNYATCIP TNHNQSPQTV

**Ph2**

LFSPMNVISP QQQQNLLQSM AAAAQQQQLT QQQQQFNQQQ QQQLTQQQQQ
LTAALAKVGV DAQGKLAQKV VQKVTTTSSA VQAATGPGST GSTQTQQVQQ
VQQQQQQTTQ TTQQCVQVSQ STLPVGVGGQ SVQTAQLLNA GQAQQMQIPW
FLQNAAGLQP FGPNQIILRN QPDGTQGMFI QQQPATQTLQ TQQNQIIQCN
VTQTPTKART QLDALAPKQQ QQQQQVGTTN QTQQQQLAVA TAQLQQQQQQ
LTAAALQRPG APVMPHNGTQ VRPASSVSTQ TAQNQSLLKA KMRNKQQPVR
PALATLKTEI GQVAGQNKVV GHLTTVQQQQ QATNLQQVVN AAGNKMVVMS
TTGTPITLQN GQTLHAATAA GVDKQQQQLQ LFQKQQILQQ QQMLQQQIAA
IQMQQQQAAV QAQQQQQQQV SQQQQVNAQQ QQAVAQQQQA VAQAQQQQRE
QQQQVAQAQA QHQQALANAT QQILQVAPNQ FITSHQQQQQ QQLHNQLIQQ
QLQQQAQAQV QAQVQAQAQQ QQQQREQQQN IIQQIVVQQS GATSQQTSQQ

**Ph3**

QQHHQSGQLQ LSSVPFSVSS STTPAGIATS SALQAALSAS GAIFQTAKPG
TCSSSSPTSS VVTITNQSST PLVTSSTVAS IQQAQTQSAQ VHQHQQLISA
TIAGGTQQQP QGPPSLTPTT NPILAMTSMM NATVGHLSTA PPVTVSVTST
AVTSSPGQLV LLSTASSGGG GSIPATPTKE TPSKGPTATL VPIGSPKTPV
SGKDTCTTPK SSTPATVSAS VEASSSTGEA LSNGDASDRS STPSKGATTP
TSKQSNAAVQ PPSSTTPNSV SGKEEPKLAT CGSLTSATST STTTTITNGI
GVARTTASTA VSTASTTTTS SGTFITSCTS TTTTTTSSIS NGSKDLPK

**Figure S2 Complete sequences of Ph IDRs.** Sequences are coloured based on amino acid type for Ph1(A), Ph2 (B), and Ph3(C). Figures were generated using Local Cider (http://pappulab.wustl.edu/CIDER/analysis/).

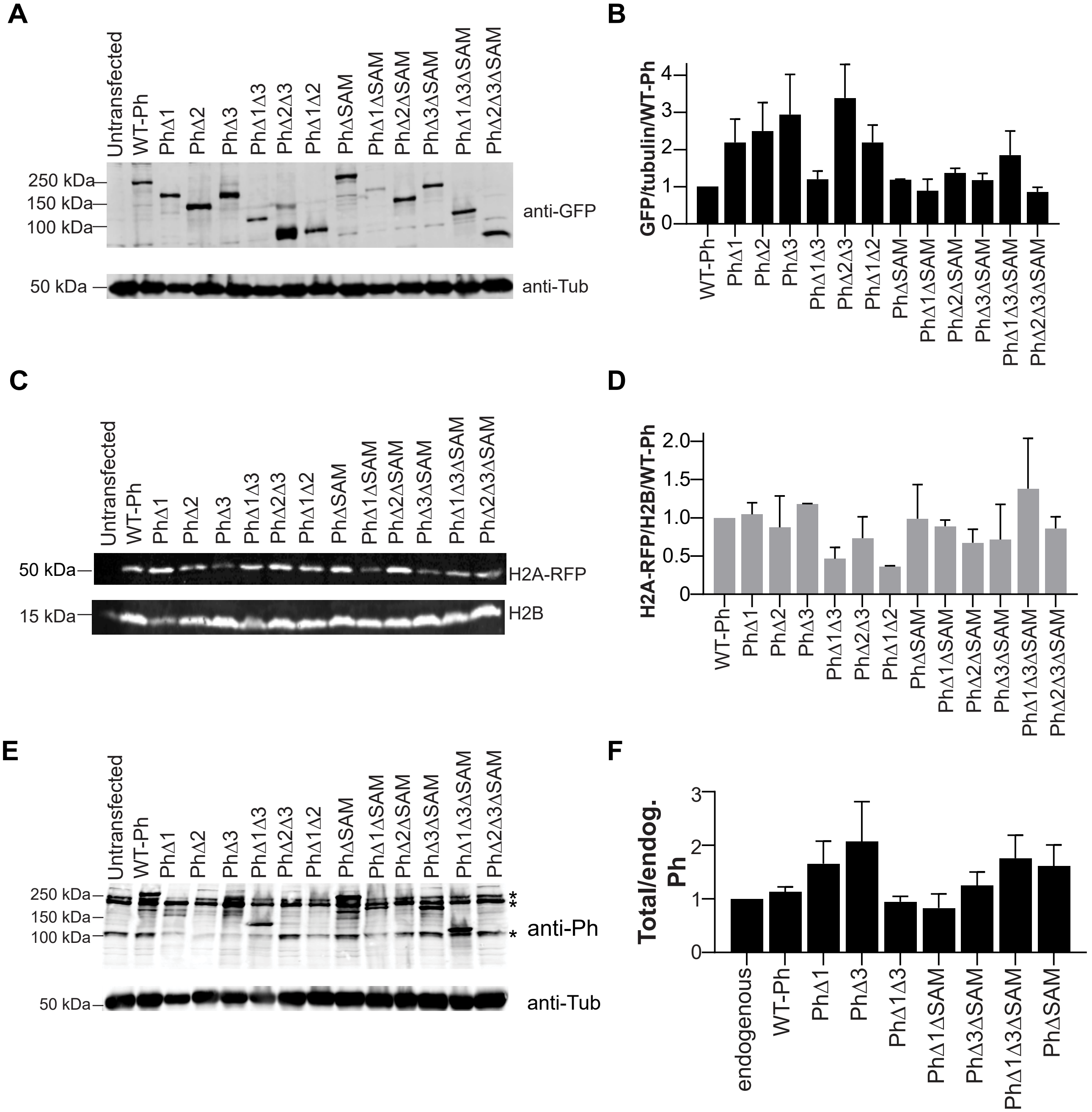
A
Untransfected
WT-Ph
PhΔ1
PhΔ2
PhΔ3
PhΔ1Δ3
PhΔ2Δ3
PhΔ1Δ2
PhΔSAM
PhΔ1ΔSAM
PhΔ2ΔSAM
PhΔ3ΔSAM
PhΔ1Δ3ΔSAM
PhΔ2Δ3ΔSAM
250 kDa
150 kDa
100 kDa
anti-GFP
50 kDa
anti-Tub
B
GFP/tubulin/WT-Ph
0
1
2
3
4
WT-Ph
PhΔ1
PhΔ2
PhΔ3
PhΔ1Δ3
PhΔ2Δ3
PhΔ1Δ2
PhΔSAM
PhΔ1ΔSAM
PhΔ2ΔSAM
PhΔ3ΔSAM
PhΔ1Δ3ΔSAM
PhΔ2Δ3ΔSAM
C
Untransfected
WT-Ph
PhΔ1
PhΔ2
PhΔ3
PhΔ1Δ3
PhΔ2Δ3
PhΔ1Δ2
PhΔSAM
PhΔ1ΔSAM
PhΔ2ΔSAM
PhΔ3ΔSAM
PhΔ1Δ3ΔSAM
PhΔ2Δ3ΔSAM
50 kDa
H2A-RFP
15 kDa
H2B
D
H2A-RFP/H2B/WT-Ph
0
0.5
1.0
1.5
2.0
WT-Ph
PhΔ1
PhΔ2
PhΔ3
PhΔ1Δ3
PhΔ2Δ3
PhΔ1Δ2
PhΔSAM
PhΔ1ΔSAM
PhΔ2ΔSAM
PhΔ3ΔSAM
PhΔ1Δ3ΔSAM
PhΔ2Δ3ΔSAM
E
Untransfected
WT-Ph
PhΔ1
PhΔ2
PhΔ3
PhΔ1Δ3
PhΔ2Δ3
PhΔ1Δ2
PhΔSAM
PhΔ1ΔSAM
PhΔ2ΔSAM
PhΔ3ΔSAM
PhΔ1Δ3ΔSAM
PhΔ2Δ3ΔSAM
250 kDa
150 kDa
100 kDa
**
*
anti-Ph
50 kDa
anti-Tub
F
Total/endog. Ph
0
1
2
3
endogenous
WT-Ph
PhΔ1
PhΔ3
PhΔ1Δ3
PhΔ1ΔSAM
PhΔ3ΔSAM
PhΔ1Δ3ΔSAM
PhΔSAM

**Figure S3 Transfected proteins are (over)expressed as full-length proteins. (A)** Representative Western blots of cells transfected with Venus-Ph proteins. **(B)** Quantification of Venus-Ph levels. Anti-GFP signal was normalized to Tubulin signal and then to WT-Ph from the same experiment. **(C)** H2Av-RFP, detected with anti-RFP, to measure transfection efficiency. **(D)** Quantification of H2Av-RFP, which was normalized to H2B and then to signal from WT-Ph from the same experiment. **(E)** Representative anti-Ph Western blot to measure total Ph levels (endogenous + transfected). Asterisks next to blots indicate the position of endogenous Ph bands. Note that the antibody epitope is in Ph2 so that proteins lacking this region are not detected. Venus-PhΔ1 and Venus-PhΔ3 migrate similarly to Ph-d. **(F)** Quantification of anti-Ph Western blots. Ph signals were normalized to tubulin, and then normalized to the signal from untransfected cells. Bars show the mean, and error bars the SEM. n=3 (B, D); n=2 (F).

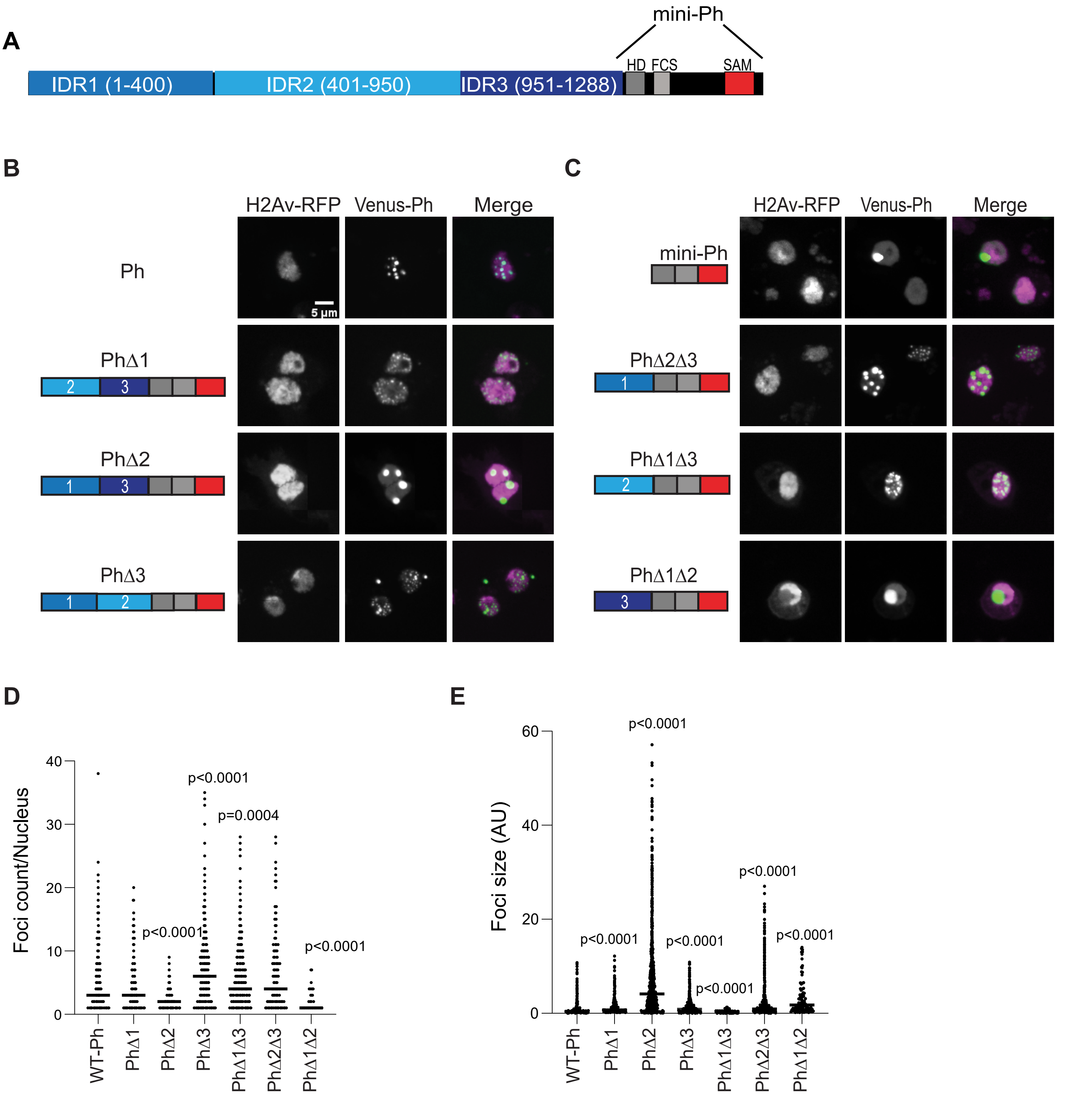
A
mini-Ph
HD FCS SAM
IDR1 (1-400)
IDR2 (401-950)
IDR3 (951-1288)
B
H2Av-RFP
Venus-Ph
Merge
Ph
5 μm
PhΔ1
PhΔ2
PhΔ3
C
mini-Ph
PhΔ2Δ3
PhΔ1Δ3
PhΔ1Δ2
D
Foci count/Nucleus
p<0.0001
p=0.0004
p<0.0001
p<0.0001
WT-Ph
PhΔ1
PhΔ2
PhΔ3
PhΔ1Δ3
PhΔ2Δ3
PhΔ1Δ2
E
Foci size (AU)
p<0.0001
p<0.0001
p<0.0001
p<0.0001
p<0.0001
p<0.0001
WT-Ph
PhΔ1
PhΔ2
PhΔ3
PhΔ1Δ3
PhΔ2Δ3
PhΔ1Δ2

**Figure S4 Summary of effects of Ph IDRs on condensate formation.** (**A**) Schematic of Ph, indicating IDR boundaries. **(B, C**) Representative images showing effects of removing one (B), or two (C) IDRs. (D, E) Graphs of number of foci per nucleus (D) and size (E). Data are taken from Fig. 3-5, but repeated to allow full comparison. See main figure legends for n-values.

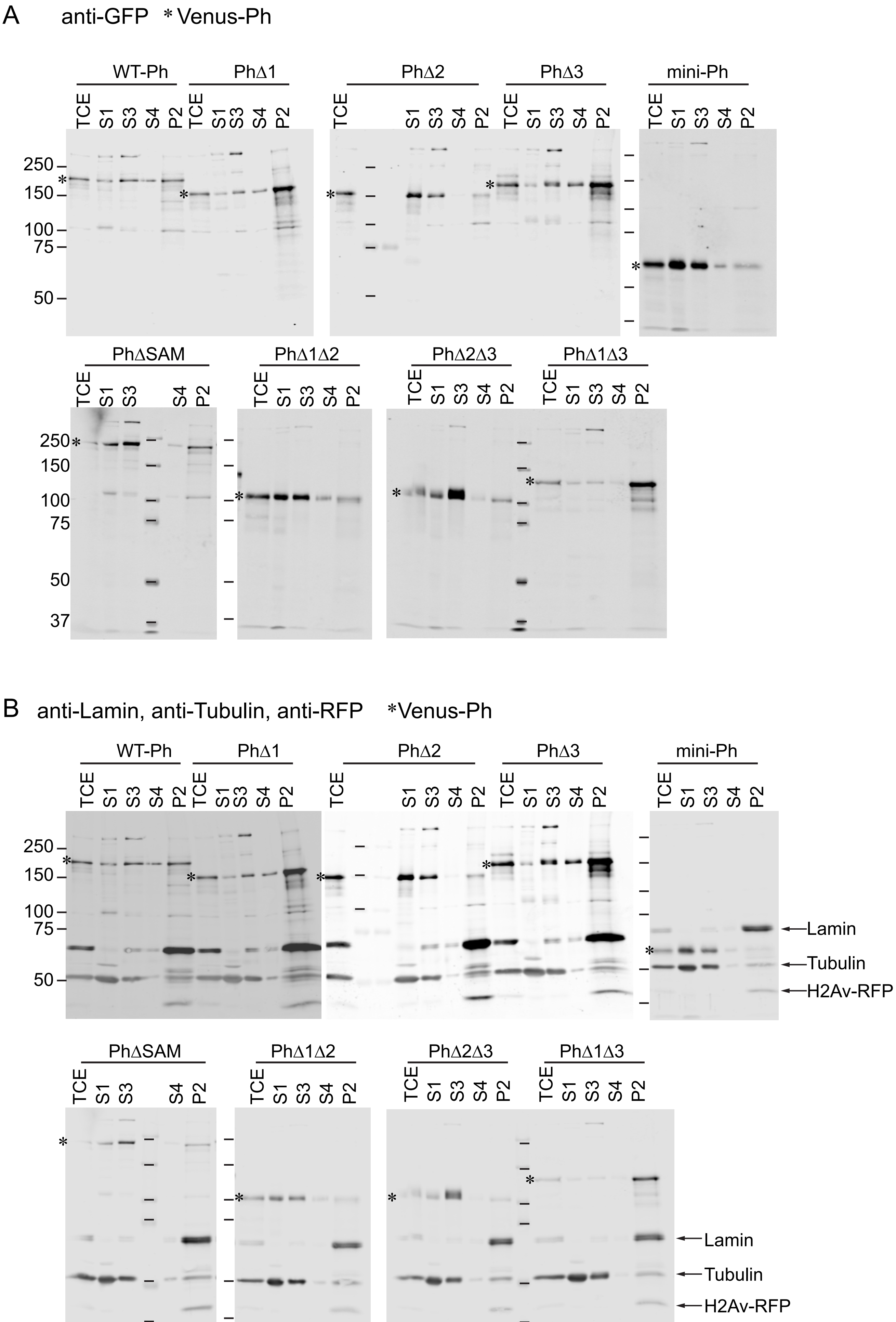
A
anti-GFP * Venus-Ph
WT-Ph
PhΔ1
PhΔ2
PhΔ3
mini-Ph
PhΔSAM
PhΔ1Δ2
PhΔ2Δ3
PhΔ1Δ3
TCE
S1
S3
S4
P2
250
150
100
75
50
37
B
anti-Lamin, anti-Tubulin, anti-RFP *Venus-Ph
Lamin
Tubulin
H2Av-RFP

**Figure S5 Full gels of Western blots of cell fractionation.** (**A**) Full blots of anti-GFP Western blots used to quantify Venus-Ph proteins in different cellular fractions (Fig. 6). TCE=total cell extract. Half as much TCE and twice as much P2 were loaded to avoid signal saturation. Asterisks indicate position of each Venus-Ph protein. (**B**). The same blots as in A were reprobed with anti-Lamin, anti-Tubulin, and anti-RFP (to detect H2Av-RFP). The anti-GFP signal is still visible, and indicated with an asterisk. Anti-Lamin and anti-Tubulin were probed with different secondaries, but the images were merged.

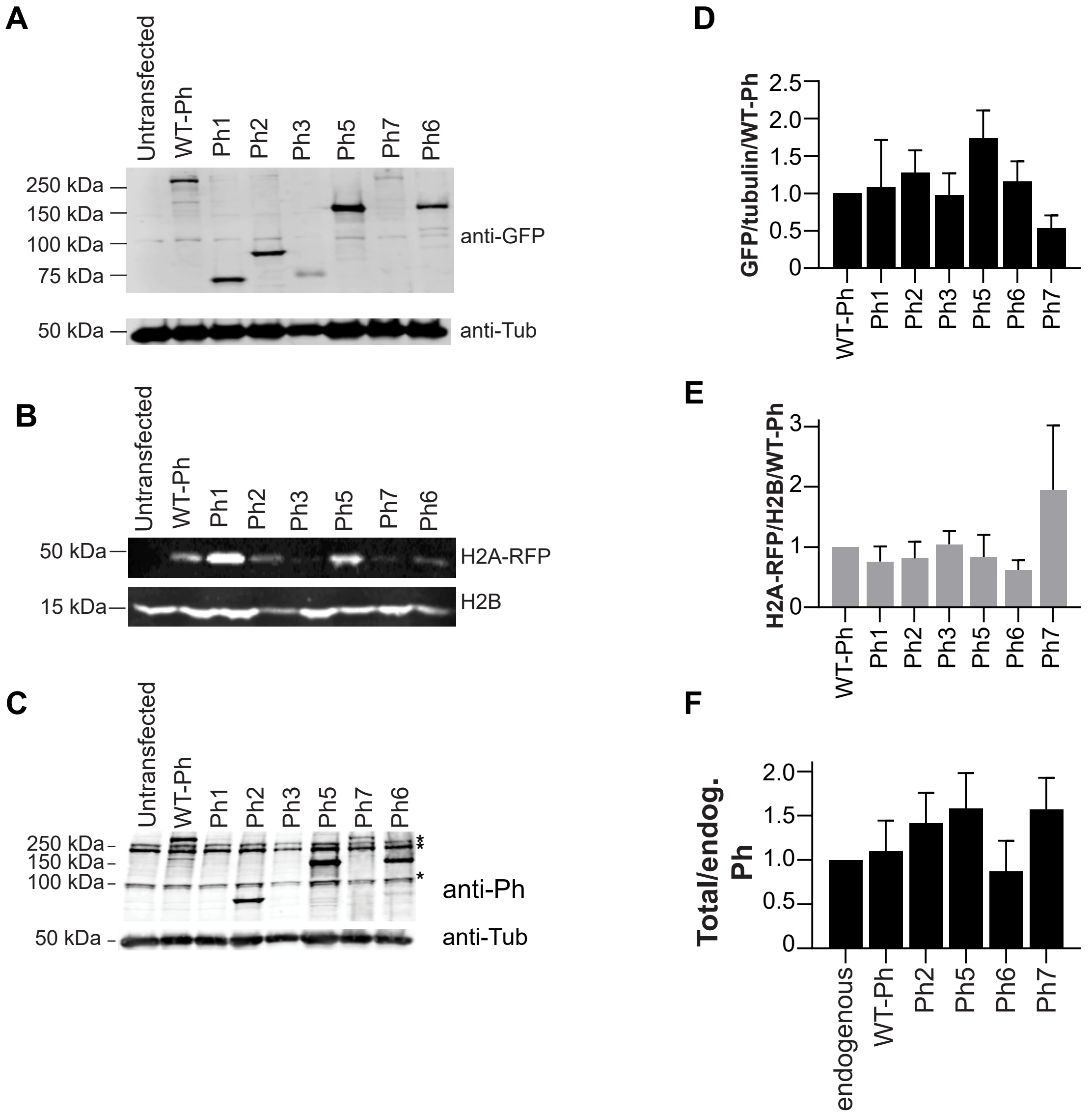
A
Untransfected
WT-Ph
Ph1
Ph2
Ph3
Ph5
Ph7
Ph6
250 kDa
150 kDa
100 kDa
75 kDa
anti-GFP
50 kDa
anti-Tub
B
Untransfected
WT-Ph
Ph1
Ph2
Ph3
Ph5
Ph7
Ph6
50 kDa
H2A-RFP
15 kDa
H2B
C
Untransfected
WT-Ph
Ph1
Ph2
Ph3
Ph5
Ph7
Ph6
250 kDa
150 kDa
100 kDa
anti-Ph
50 kDa
anti-Tub
D
GFP/tubulin/WT-Ph
2.5
2.0
1.5
1.0
0.5
0
WT-Ph
Ph1
Ph2
Ph3
Ph5
Ph6
Ph7
E
H2A-RFP/H2B/WT-Ph
3
2
1
0
WT-Ph
Ph1
Ph2
Ph3
Ph5
Ph6
Ph7
F
Total/endog. Ph
2.0
1.5
1.0
0.5
0
endogenous
WT-Ph
Ph2
Ph5
Ph6
Ph7

**Figure S6 Quantification of Ph IDR expression levels in transfected cells. (A-C)** Representative Western blots of cells transfected with Venus-Ph IDR constructs. anti-GFP is used to quantify Venus (A) and anti-RFP to quantify H2Av-RFP (B) (to measure transfection efficiency). anti-Ph (C) indicates the amount of IDR+ Ph in cells. Because the antigen for the anti-Ph antibody is in Ph2, only proteins containing this region can be measured (Ph2, Ph5, Ph6, Ph7). **(D-E)** Quantification of Western blot signals. D and F are normalized to tubulin, and E to H2B. Signals are then normalized to WT-Ph from the same experiment (D, E), or untransfected (F) (endogenous Ph levels). Bars are mean and error bars are SEM. n=3 (D); n=2 (E, F).
